# scCDC: a computational method for gene-specific contamination detection and correction in single-cell and single-nucleus RNA-seq data

**DOI:** 10.1101/2022.11.24.517598

**Authors:** Weijian Wang, Yihui Cen, Zezhen Lu, Yueqing Xu, Tianyi Sun, Ying Xiao, Wanlu Liu, Jingyi Jessica Li, Chaochen Wang

## Abstract

In droplet-based single-cell RNA-seq (scRNA-seq) and single-nucleus RNA-seq (snRNA-seq) assays, systematic contamination of ambient RNA molecules biases the estimation of genuine transcriptional levels. To correct the contamination, several computational methods have been developed. However, these methods do not distinguish the contamination-causing genes and thus either under- or over-corrected the contamination in our in-house snRNA-seq data of virgin and lactating mammary glands. Hence, we developed scCDC as the first method that specifically detects the contamination-causing genes and only corrects the expression counts of these genes. Benchmarked against existing methods on synthetic and real scRNA-seq and snRNA-seq datasets, scCDC achieved the best contamination correction accuracy with minimal data alteration. Moreover, scCDC applies to processed scRNA-seq and snRNA-seq data with empty droplets removed. In conclusion, scCDC is a flexible, accurate decontamination method that detects the contamination-causing genes, corrects the contamination, and avoids the over-correction of other genes.

## Background

Single-cell RNA-seq (scRNA-seq) is a widely-used technique for studying cell heterogeneity in organs. Various studies and large databases, such as the human cell atlas, have taken advantage of scRNA-seq, especially droplet-based platforms such as Chromium X, BD Rhapsody, and inDrop [1-3]. Droplet-based scRNA-seq requires every cell to be sealed with a barcoded bead in a droplet so that the cell’s mRNAs can be labeled by the specific barcode. However, ambient RNA contamination is ubiquitous [4-7]: ambient RNA molecules in the solution would cause systematic contamination by inflating the estimation of endogenous genes’ expression levels in cells, thus impeding the identification of cell-type marker genes. In parallel to scRNA-seq, single-nucleus RNA-seq (snRNA-seq) has been developed to investigate the cells that are too fragile or difficult to dissociate into single cells [8, 9]. Yet, ambient RNA contamination is likely more common in snRNA-seq than in scRNA-seq because the nuclei extraction procedure would cause many RNAs in the cytoplasm to be released into the solution.

Various experimental and computational strategies have been developed to correct the contamination in scRNA-seq and snRNA-seq data. Sanchez et al. developed an experimental approach that uses spike-in cells as a reference to correct the contamination [6]. However, this approach complicates the experimental procedure and has not been integrated into common commercial platforms. Several computational methods have been developed, including SoupX [5], CellBender [10, 11], and scAR [10, 11], whose common idea is first estimating the distribution of ambient RNA levels from empty droplets and then using the estimated distribution to correct the gene expression levels in cells. Since SoupX, CellBender, and scAR require empty-droplet data, they are inapplicable to processed data in which empty droplets have been removed. Although another computational method, DecontX [4], does not require empty-droplet data, it and the three other methods do not distinguish the contamination-causing genes but alter all genes’ expression levels, possibly leading to over-correction.

In this study, we performed snRNA-seq assays in mouse mammary glands at the virgin and lactation stages. In our snRNA-seq datasets, we observed sample-specific contamination by ambient RNAs. To correct the contamination, we applied the existing computational methods mentioned above but found that DecontX and CellBender had under-correction, while SoupX and scAR over-corrected many genes, including housekeeping genes (Figure 1D&E).

**Figure 1.**
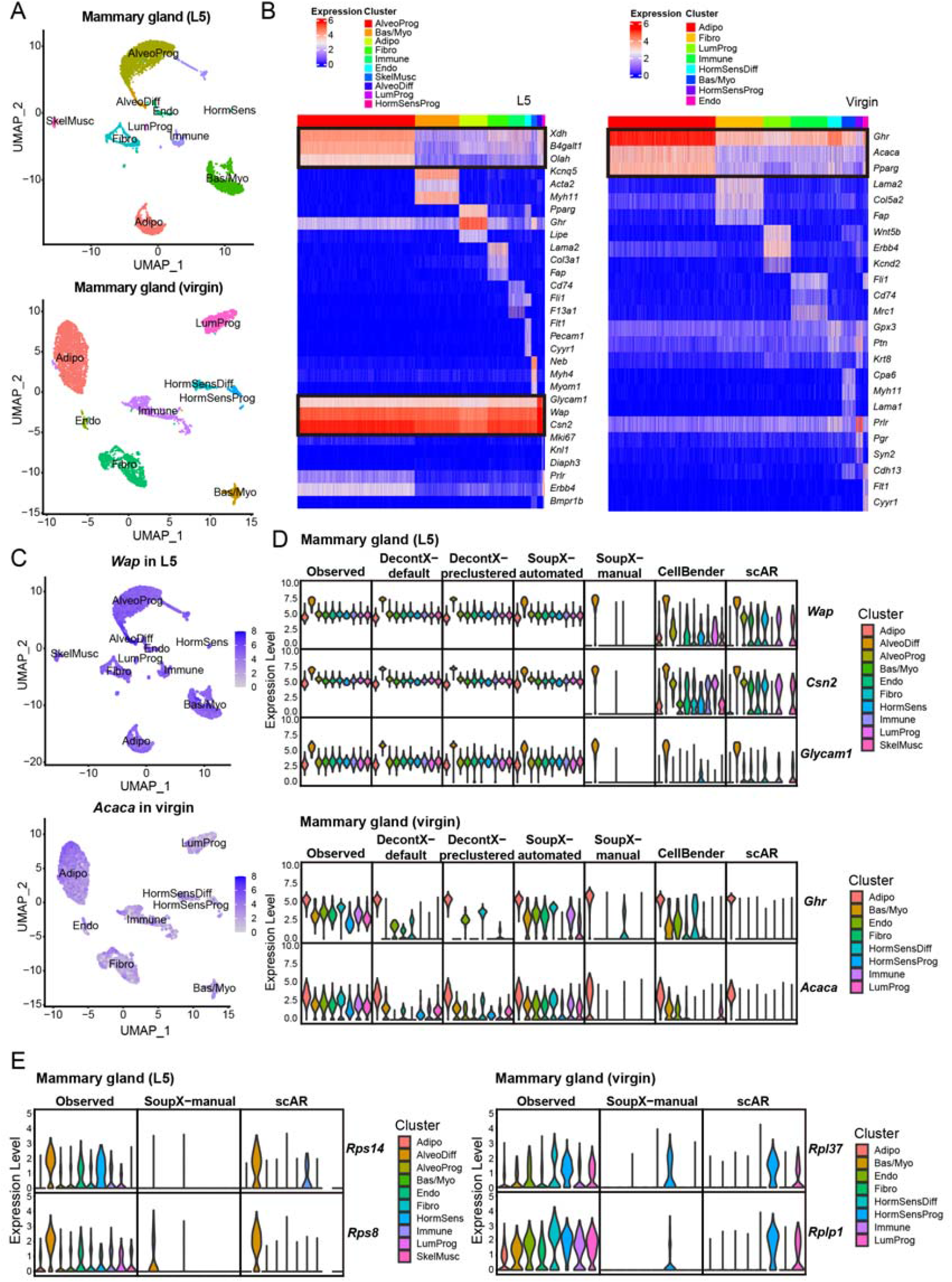
Performance evaluation of existing tools on correcting contaminative mammary gland snRNA-Seq data. (A) The cell clusters identified in L5 and virgin mammary gland datasets are shown in UMAP plots. (B) Heatmap of the expression of selected marker genes in L5 and virgin mammary gland datasets. Notably, highlighted genes supposed to express exclusively in a cluster are widely detected in all the cells. (C) The expression of *Wap* and *Acaca* in the nucleus are shown in UMAP plots. (D-E) The violin plots show the normalized expression levels of the selected marker genes (D) and housekeeping genes (E) before and after correction using the indicated methods by the default Seurat (V3). Adipo, adipocytes; AlveoProg, alveolar progenitors; AlveoDiff, differentiated alveolar cells; Bas/Myo, basal cells/myoepithelial cells; Endo, endothelial cells; Fibro, fibroblasts; HormSens, hormone sensing cells; HormSensDiff, differentiated hormone sensing cells; HormSensProg, hormone sensing progenitors; Immune, immune cells; LumProg, luminal progenitors; SkelMusc, skeleton muscle cells.

Motivated by the limitations of existing computational methods, we developed scCDC (single-cell Contamination Detection and Correction), which first detects the “contamination-causing genes,” which encode the most abundant ambient RNAs, and then only corrects these genes’ measured expression levels. Benchmarked against DecontX, SoupX, CellBender, and scAR, scCDC achieved the best contamination correction accuracy with minimal data alteration. Not requiring empty-droplet data, scCDC applies to all processed scRNA-seq and snRNA-seq datasets in public repositories. We further showed that scCDC improved the accuracy of identifying cell-type marker genes and constructing gene co-expression networks.

## Results

### Ambient RNAs contaminated snRNA-seq data of mouse mammary glands

The mammary gland is a unique mammalian organ whose sole function is to feed the young. Hence, the mammary gland undergoes dramatic developmental changes during pregnancy and lactation. To investigate mammary gland development, several studies have performed scRNA-seq on epithelial cells of mammary glands [12-15]. However, the development of mammary glands also requires the interplay between epithelial cells and the cells in the niche, including adipocytes, fibroblasts, and immune cells [16-18].

Instead of scRNA-seq, we employed snRNA-seq to profile a complete cellular map in virgin and lactating (lactation day 5, denoted by L5) mouse mammary glands. In addition to epithelial cells and subsets of luminal and basal cells, we successfully identified adipocytes, fibroblasts, and immune cells, which had not been efficiently captured by previous scRNA-seq studies (Figure 1A). However, we found several well-known cell-type marker genes unexpectedly detected in nearly all cell types. For example, the genes *Wap* and *Csn2*, which encode whey acidic protein and casein protein respectively, should be expressed exclusively in the differentiated alveolar epithelial cells (AlveoDiff) during lactation; the gene *Acaca*, which encodes the acetyl-CoA carboxylase for fatty acid synthesis, is expected to be expressed exclusively in adipocytes (Adipo). Surprisingly, however, these genes’ mRNAs were also detected in nearly all the other cell types. Similarly, AlveoDiff marker *Glycam1* and Adipo marker *Ghr* were detected globally in lactating and virgin datasets, respectively (Figure 1 B&C). These data suggested the presence of systematic contamination by ambient RNAs.

### Performance evaluation of existing correction methods on mouse mammary gland snRNA-seq datasets

The four aforementioned computational methods—DecontX, SoupX, CellBender, and scAR—were developed to correct contaminated scRNA-seq and snRNA-seq data [4-6, 10, 11]. Here, we benchmarked the performance of these methods in correcting our in-house snRNA-seq data of mouse mammary glands.

Applied to the lactating dataset, DecontX barely removed any contamination of AlveoDiff markers *Wap, Csn2*, and *Glycam1* in both the “default” mode (DecontX-default) and the “pre-clustered” mode that takes user-specified cell clusters (DecontX-pre-clustered). Similarly, SoupX failed to correct the three genes’ contamination in the “automated” mode (SoupX-automated), and SoupX “manual” mode (SoupX-manual, which takes user-defined contamination-causing genes) only achieved a reasonable correction performance. Moreover, CellBender and scAR under-corrected the three genes’ contamination (Figure 1D, upper panel).

Applied to the virgin dataset, only scAR successfully corrected the contamination of Adipo markers *Ghr* and *Acaca*. Specifically, SoupX-automated failed to correct these two genes’ contamination; DecontX-default, DecontX-pre-clustered, and CellBender all under-corrected these two genes’ contamination; SoupX-manual under-corrected *Ghr*’s contamination (Figure 1D, lower panel).

Since DecontX, SoupX, CellBender, and scAR alter all genes’ counts, we also checked how they altered the counts of the genes other than the above cell-type marker genes. Although SoupX-manual and scAR had less of an under-correction issue for cell-type marker genes, they undesirably removed the counts of housekeeping genes, such as *Rps14, Rps8, Rpl37*, and *Rplp1*, in multiple cell types (Figure 1E). Examining the counts of 66 housekeeping genes before and after each method’s correction, we found that SoupX-manual and scAR undesirably removed the counts of many housekeeping genes in more than 95% of cells (Supplementary Figure 1). These results revealed the over-correction issue of SoupX-manual and scAR.

Taken together, our benchmark results show that DecontX, SoupX-automated, and CellBender under-corrected the contamination-causing genes (usually cell-type marker genes), while SoupX-manual and scAR over-corrected the uncontaminated genes, including house-keeping genes.

### Overview of scCDC

The existing correction methods’ limitations, in particular, SoupX-manual and scAR’s over-correction, motivated us to devise a strategy to identify contamination-causing genes and correct only these genes’ contamination. In contrast to the global strategy used by the existing correction methods, this gene-specific strategy can better avoid the under-correction of contamination-causing genes and the over-correction of other genes. Following this strategy, we developed a new method, scCDC, which has two functionalities: contamination detection and correction.

A contamination-causing gene has abundant ambient RNAs, so its observed count in a droplet is the sum of the counts from its endogenous and ambient RNAs (Figure 2A). scCDC is designed to identify such a gene first and then correct the gene’s observed counts. Because the gene’s ambient RNAs are abundant but less variable in droplets, the ambient RNA counts would deflate the entropy of the gene’s observed counts. Following this rationale, scCDC’s contamination detection functionality consists of three steps (Figure 2B). (Note that scCDC requires cells to be pre-clustered, an issue we discussed in the Method Appendix.) First, under the assumption that most genes produce little or no ambient RNAs (defined as “endogenous genes”), scCDC estimates the expected entropy-expression curve of endogenous genes within each cell cluster (see Methods). Second, in each cell cluster, scCDC calculates the “entropy divergence,” defined as a gene’s expected entropy (which is calculated based on the gene’s expression and the expected entropy-expression curve) minus its observed entropy, to represent the gene’s contamination level. Third, scCDC identifies the “global contamination-causing genes” (GCGs, details in Methods) as the genes with statistically significant entropy divergences in more than 80% of the cell clusters. (Note that 80% is the default value of the “restriction factor,” a tuning parameter that can be user-specified: the larger the restriction factor, the fewer GCGs scCDC identifies; we set the default restriction factor to 80% based on empirical results—details in Methods and the Method Appendix.)

**Figure 2.**
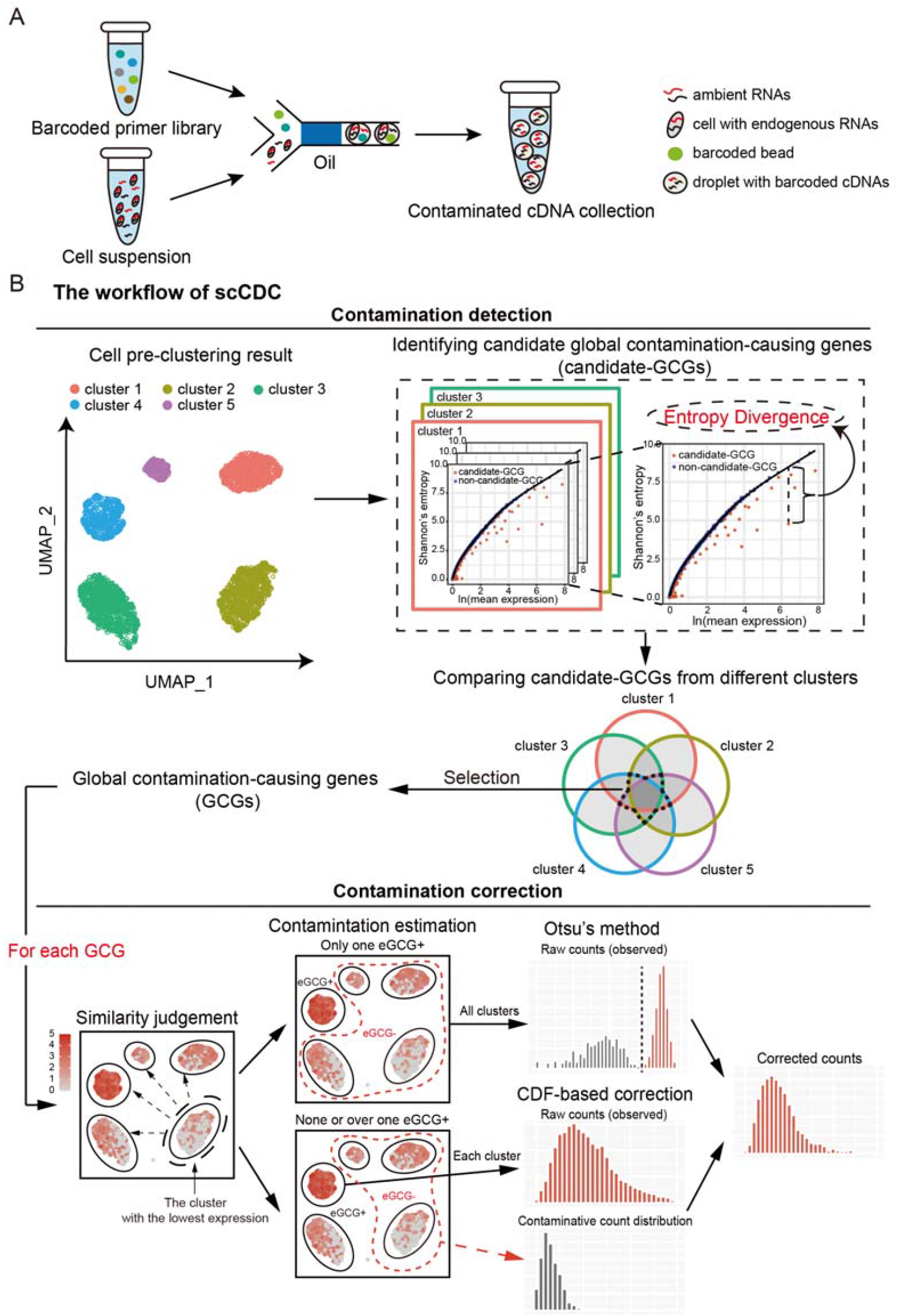
An overview of scCDC workflow. (A) The diagram of contamination shows ambient RNAs cause contaminated profiles for scRNA-Seq (or snRNA-Seq). (B) Workflow of scCDC. The theoretical entropy-expression curves of endogenous RNAs are simulated and the divergence of observed and expected entropy are calculated. Genes with significant entropy divergence were selected in each cluster and the common genes were defined as GCGs. For contamination correction, the clusters of cells do or do not express endogenous GCGs were first defined (eGCG+ and eGCG-cells). Otsu’s method and Cumulative distribution function (CDF)-based approach were used to correct the contaminated counts depending on the number of eGCG+ cell clusters (details in Methods).

After detecting the GCGs, scCDC’s contamination correction functionality corrects these GCGs’ observed counts. For each GCG, scCDC corrects the observed counts in two steps. First, scCDC finds the cell clusters in which the GCG is unlikely expressed and labels these clusters as eGCG- and the remaining clusters as eGCG+ (Figure 2B). Technically, scCDC locates the cell cluster in which the GCG has the lowest mean expression; then, scCDC groups the cell cluster with similar clusters in terms of the *Wasserstein* distance (based on the GCG’s count distribution in each cluster; details in Methods). The justification is that the GCG should have similar count distributions in the clusters where it is unexpressed because its ambient RNAs determine its count distributions in these clusters. Second, scCDC corrects the GCG’s counts using the GCG’s count distributions in the eGCG- and eGCG+ cell clusters by the *Otsu*’s method or by a cumulative distribution function (CDF)-based correction approach (Figure 2B and Methods; the *Otsu*’s method is only applicable when the GCG has only one eGCG+ cluster; Supplementary Figure 2 shows that empirically the *Otsu*’s method has better correction accuracy when it is applicable).

Table 1 compares scCDC with DecontX, SoupX, CellBender, and scAR using five criteria: (1) whether a method can work without empty-droplet data, (2) whether a method can run with CPU only, (3) whether a method corrects all genes, (4) whether a method evaluates a gene’s contamination within each cell cluster, and (5) whether a method requires preclustering. For scCDC, the answers are yes, yes, no, yes, and yes.

**Table 1.** summarizes the characteristics of scCDC in contrast to those of DecontX, SoupX, CellBender, and scAR.

|  | Only a filtered gene-by-cell matrix needed (no empty droplets) | Only CPU needed | Data correction | Contamination evaluation in individual cluster | Preclustering required |
| --- | --- | --- | --- | --- | --- |
| scCDC | √ | √ | GCGs only | √ | √ |
| DecontX-default | √ | √ | Globally | × | √ |
| DecontX-preclustered | √ | √ | Globally | × | √ |
| SoupX-automated | × | √ | Globally | × | × |
| SoupX-manual | × | √ | Globally | × | × |
| CellBender | × | × | Globally | × | × |
| scAR | × | √ | Globally | × | × |

### Simulation validated scCDC’s contamination detection and correction functionalities

To validate the performance of scCDC in a scenario with ground truths, we first simulated an uncontaminated PBMC single-cell dataset by a realistic simulator scDesign2 [19] and then artificially contaminated the data with “ambient” counts of three CD14+ monocyte marker genes *LYZ, S100A8*, and *S100A9*. As expected, the three genes’ entropy divergences increased strikingly in all cell clusters after the artificial contamination (Figure 3A). More importantly, the three genes’ entropy divergences correlated positively with the artificial contamination levels (i.e., the proportions of “ambient” counts), suggesting that the entropy divergence is a reasonable measure of the contamination level (Figure 3B). As expected, scCDC successfully identified all three genes as GCGs. These results supported scCDC’s contamination detection functionality.

**Figure 3.**
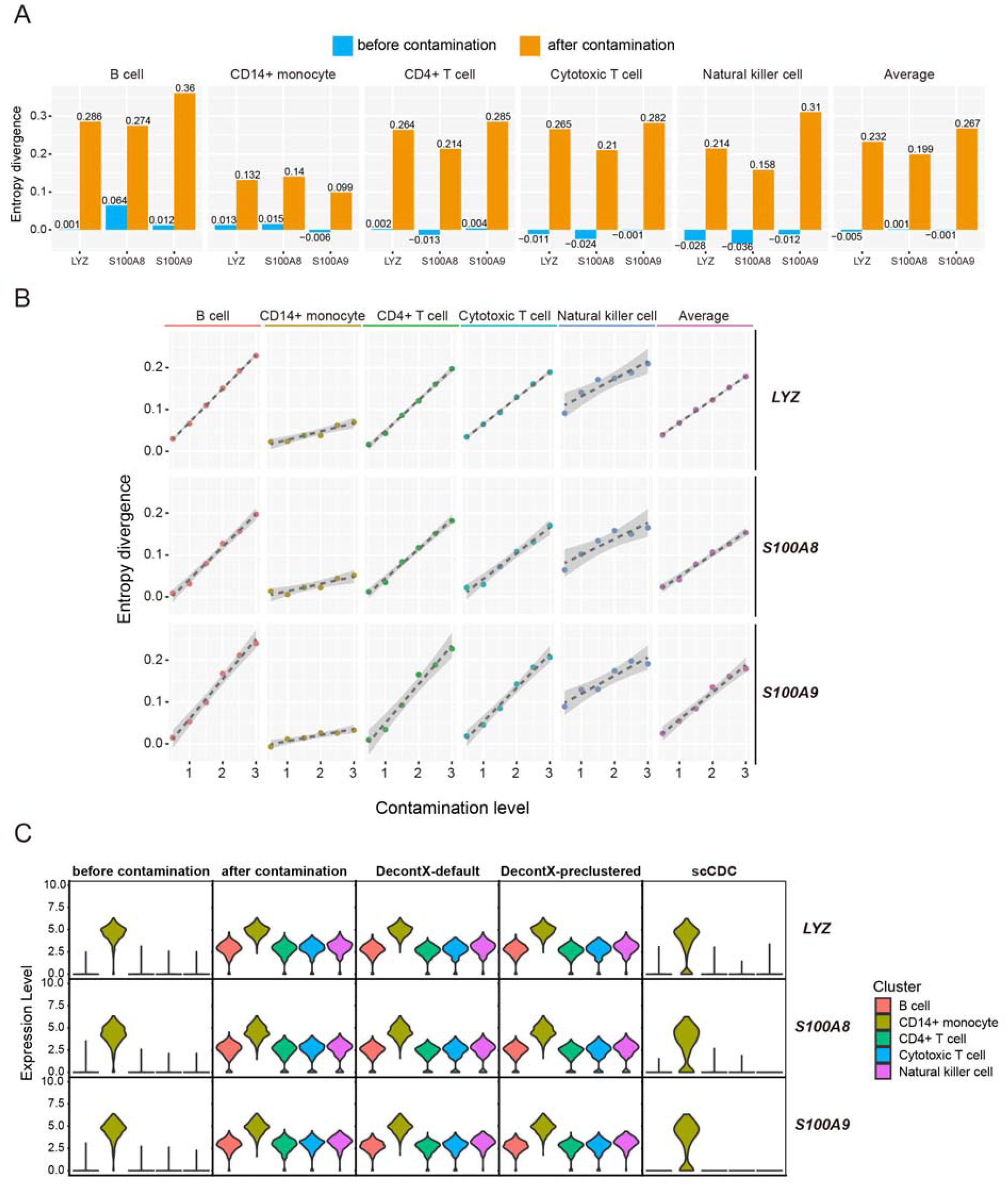
Simulation confirms the functionalities of scCDC. (A) The bar plots show the entropy divergence from the simulated curves for the three artificially contaminated genes in simulated PBMC data. (B) The scatter plots show the positive correlation between the entropy divergence and the artificial contamination level. Different contamination level was obtained by adding simulated contaminative counts with indicated mean values (0.5, 1.0, 1.5, 2.0, 2.5, 3.0). For each mean, various distribution size (1, 10, 50, 100) were employed, and the average entropy divergence were calculated and shown. (C) Benchmarking scCDC and DecontX in simulated dataset. The violin plots show the normalized expression levels of the three artificial GCGs before and after correction using the indicated methods by the default Seurat (V3).

We further validated scCDC’s contamination correction functionality on this simulated dataset and found that scCDC successfully corrected the contamination of all three GCGs (Figure 3C). We also benchmarked scCDC against DecontX (note that the other existing methods are inapplicable because they require the counts of empty droplets, which are unavailable in the simulated dataset). In contrast to scCDC, DecontX-default and DecontX-pre-clustered barely removed any contamination (Figure 3C), consistent with their performance on our in-house mouse mammary gland snRNA-seq data (Figure 1D).

### Benchmarking scCDC against existing methods on real snRNA-seq and scRNA-seq datasets

Applied to our in-house snRNA-seq datasets of mouse mammary glands, scCDC detected 32 and 38 GCGs in lactating and virgin mammary glands, respectively (Table 2). Consistent with our knowledge, AlveoDiff marker genes, including *Wap, Csn*, and *Glycam1*, were identified as GCGs in the lactating mammary glands; Adipo marker genes, such as *Acaca, Cidec*, and *Ghr*, were found as GCGs in the virgin mammary glands (Figure 4A). Examining the corresponding bulk RNA-seq data of the mammary glands, we confirmed that these identified GCGs were highly expressed in the corresponding tissues (Figure 4B), consistent with the fact they likely caused global contamination. Furthermore, scCDC successfully removed the GCGs’ contamination (Figure 4C), on a par with SoupX-manual and better than all other existing methods (Supplementary Figure 3).

**Table 2.**
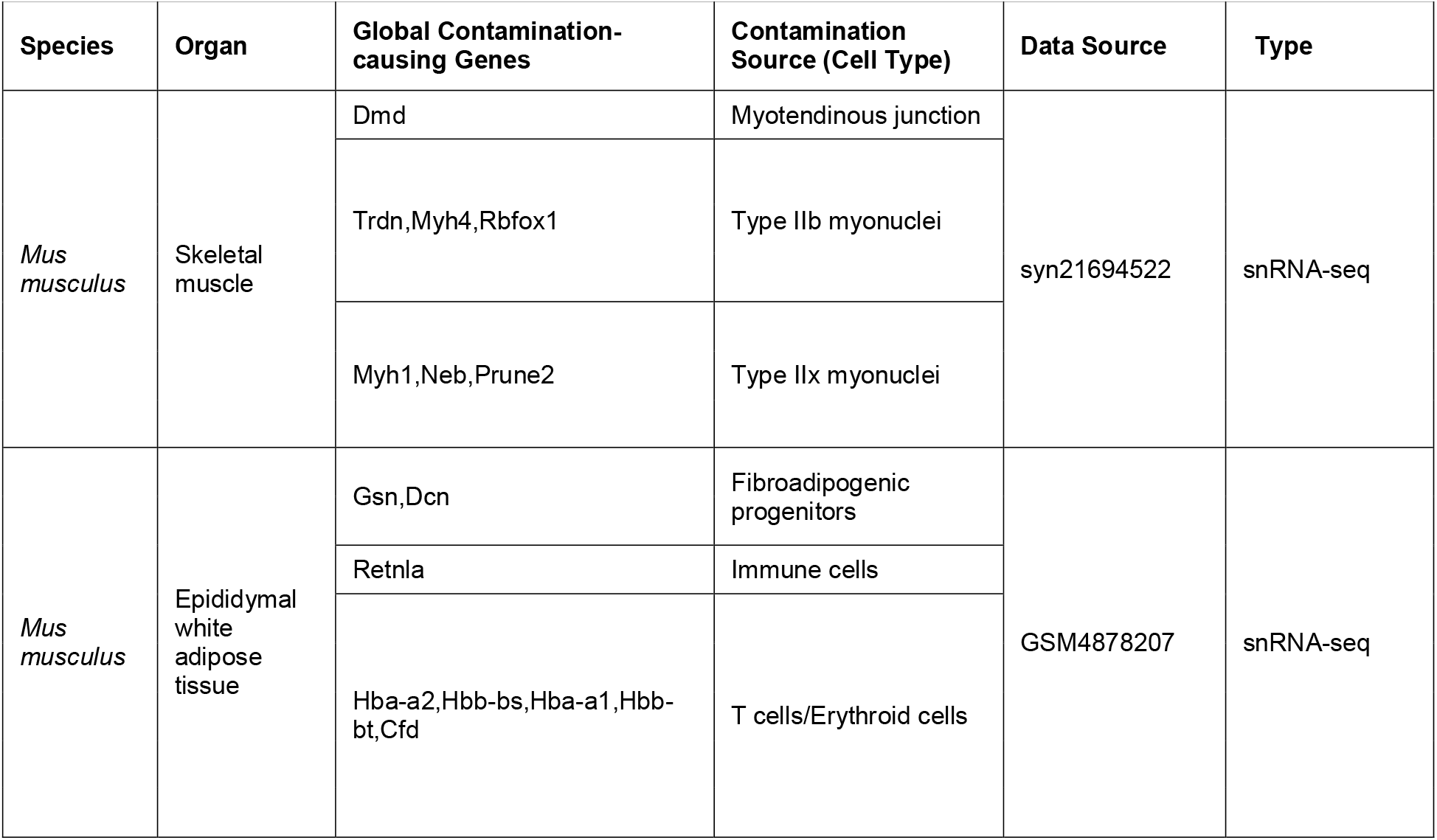
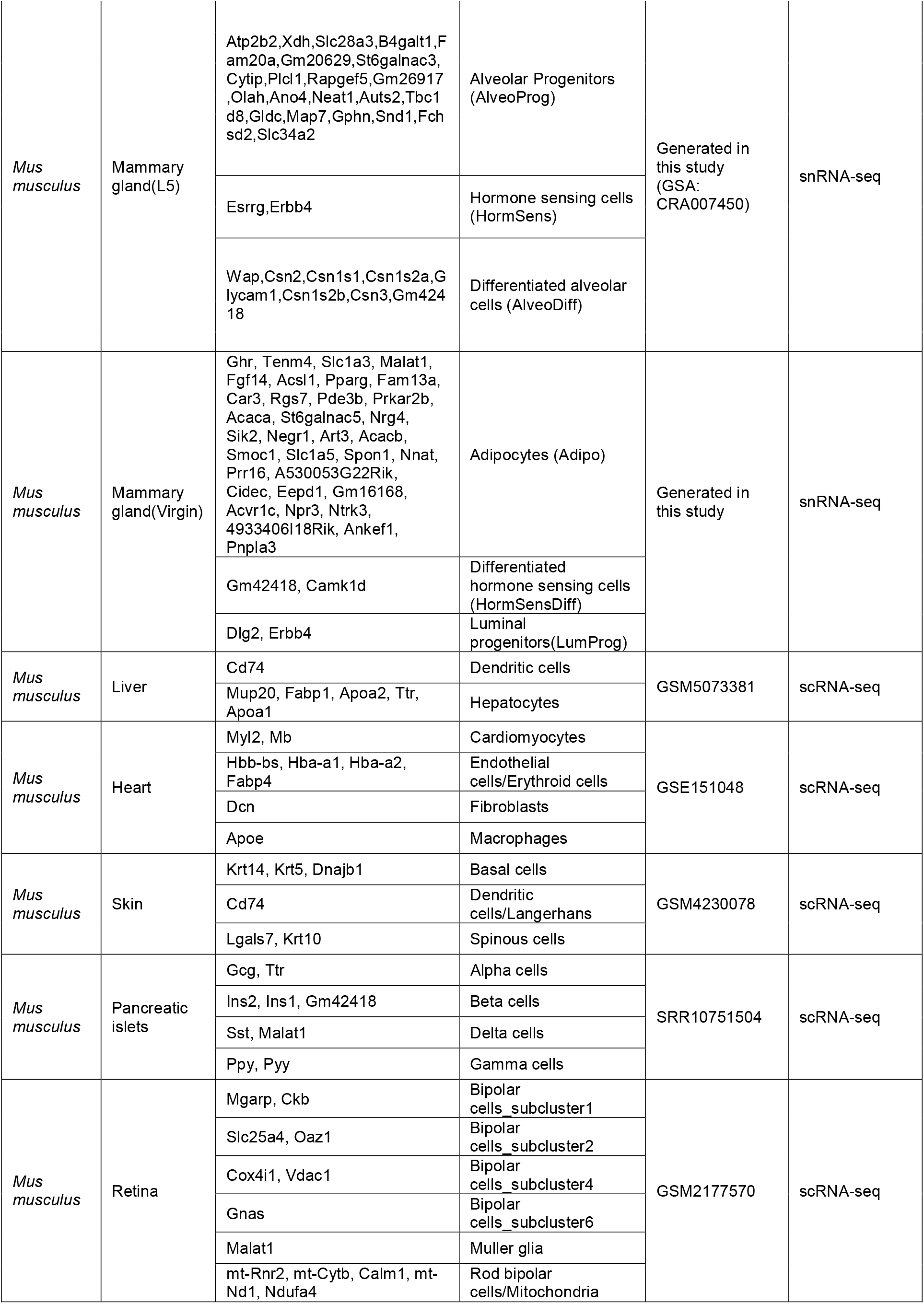
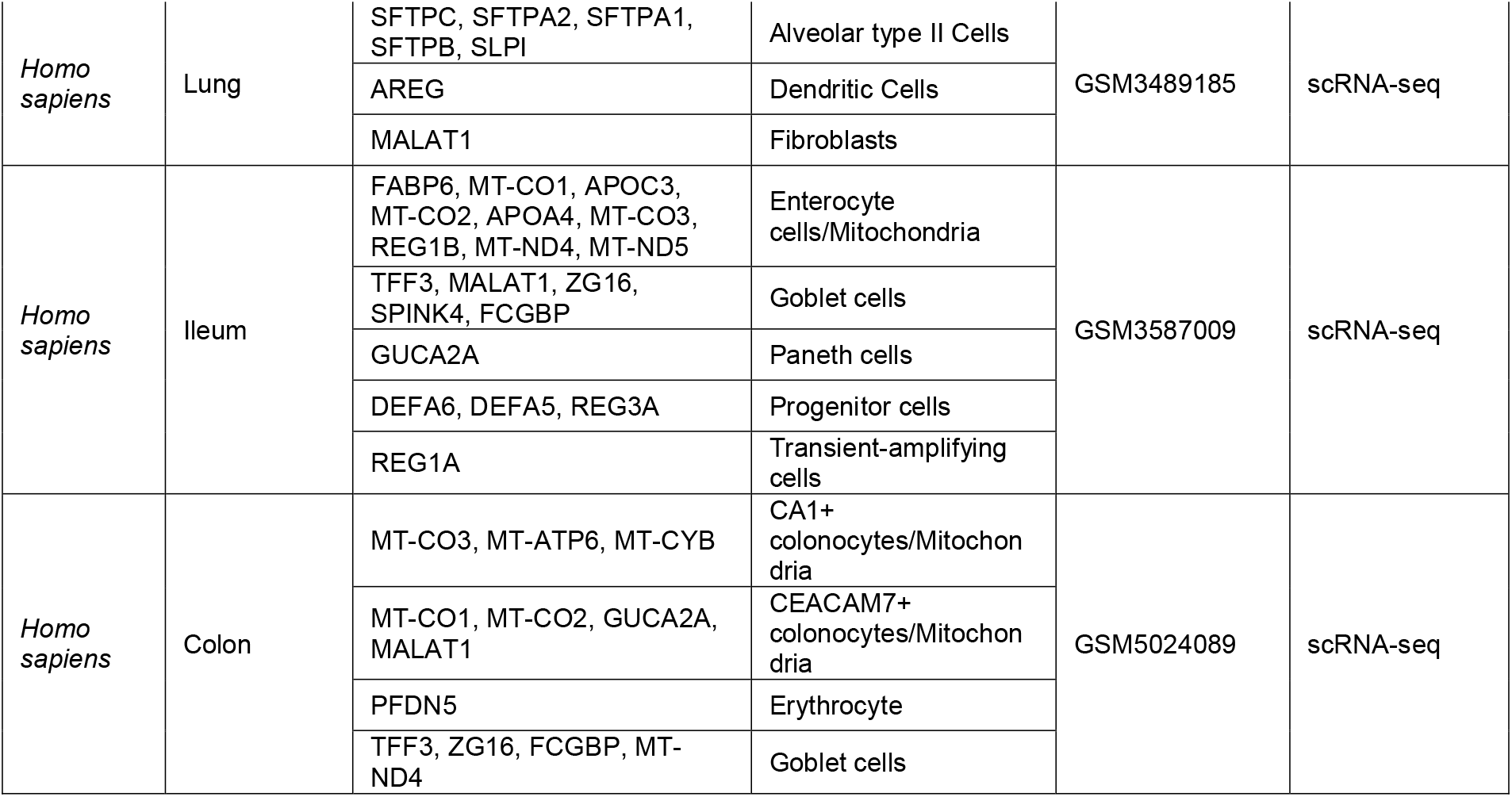
GCGs identified by scCDC in 12 snRNA-seq and scRNA-seq datasets.

**Figure 4.**
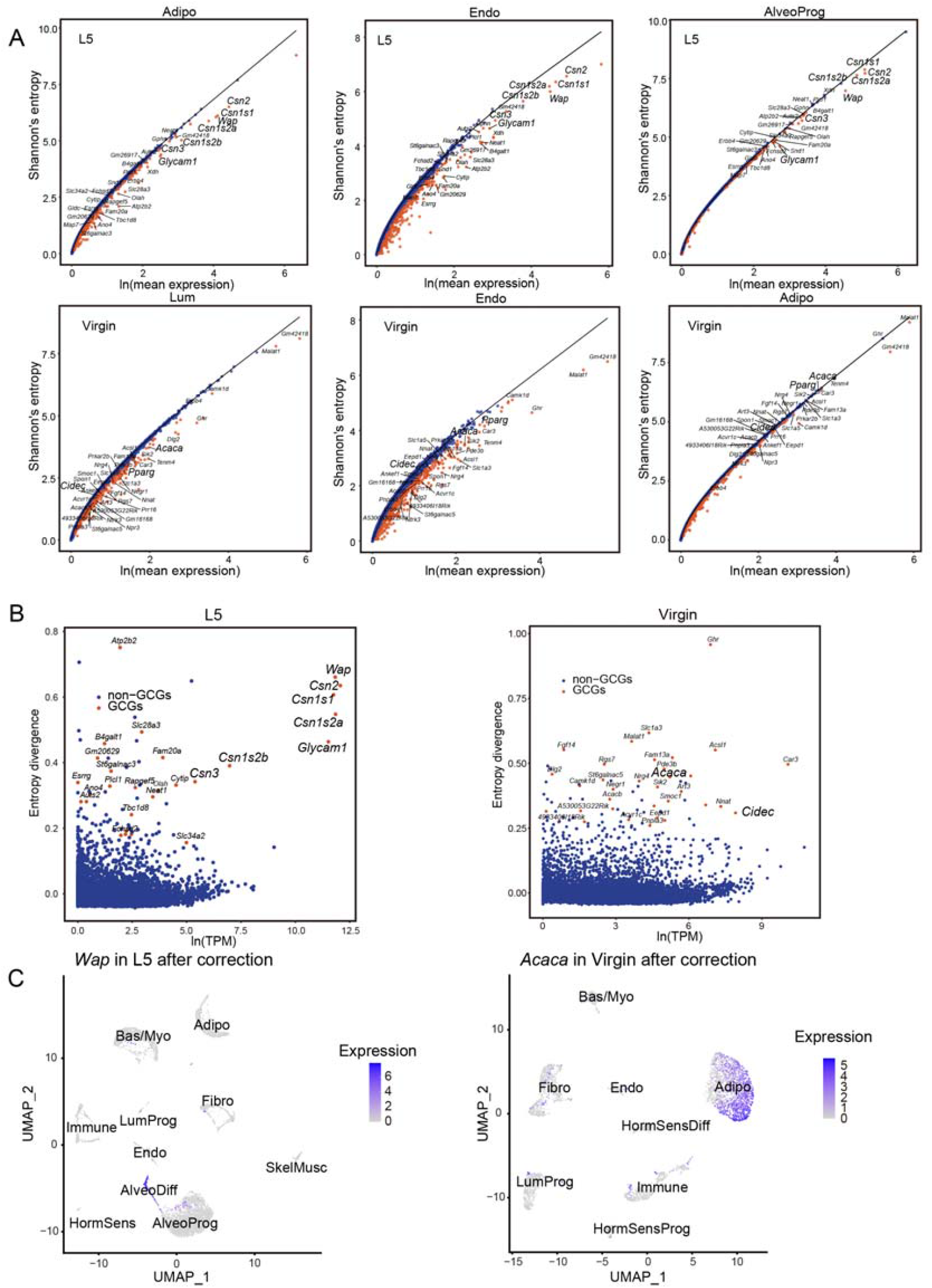
Contamination detection in snRNA-Seq datasets of mammary gland. (A) The entropy-expression curves of indicated cells. Identified GCGs were labeled on the plot. (B) Scatter plot of the relationship between the mean entropy divergence and the TPM (x-axis, log scale) of all the genes. GCGs are highlighted in red. (C) The expression of *Wap* and *Acaca* in the nucleus after correction are shown in UMAP plots.

Sanchez et al. also reported the contamination by ambient RNAs in pancreas scRNA-seq data [6]. For instance, insulin encoding genes, *Ins1* and *Ins2*, should be expressed exclusively and abundantly in beta cells, but they were unexpectedly detected in almost all cells (Supplementary Figure 4). Applying scCDC to the scRNA-seq data, we identified nine GCGs, including *Ins1* and *Ins2*, consistent with Sanchez et al.’s original finding (Table 2). Again, eight GCGs identified by scCDC were confirmed as highly expressed by bulk RNA-seq (Figure 5A; the only GCG not found in the bulk RNA-seq data was a pseudogene). Moreover, we found the GCGs’ count distributions deviated significantly from the negative-binominal (NB) distribution, i.e., the expected count distribution without contamination [19, 20]; in contrast, the housekeeping genes’ count distributions follow the NB distribution well. This contrastive result confirms the existence of contamination by the GCGs’ ambient RNAs (Figure 5B).

**Figure 5.**
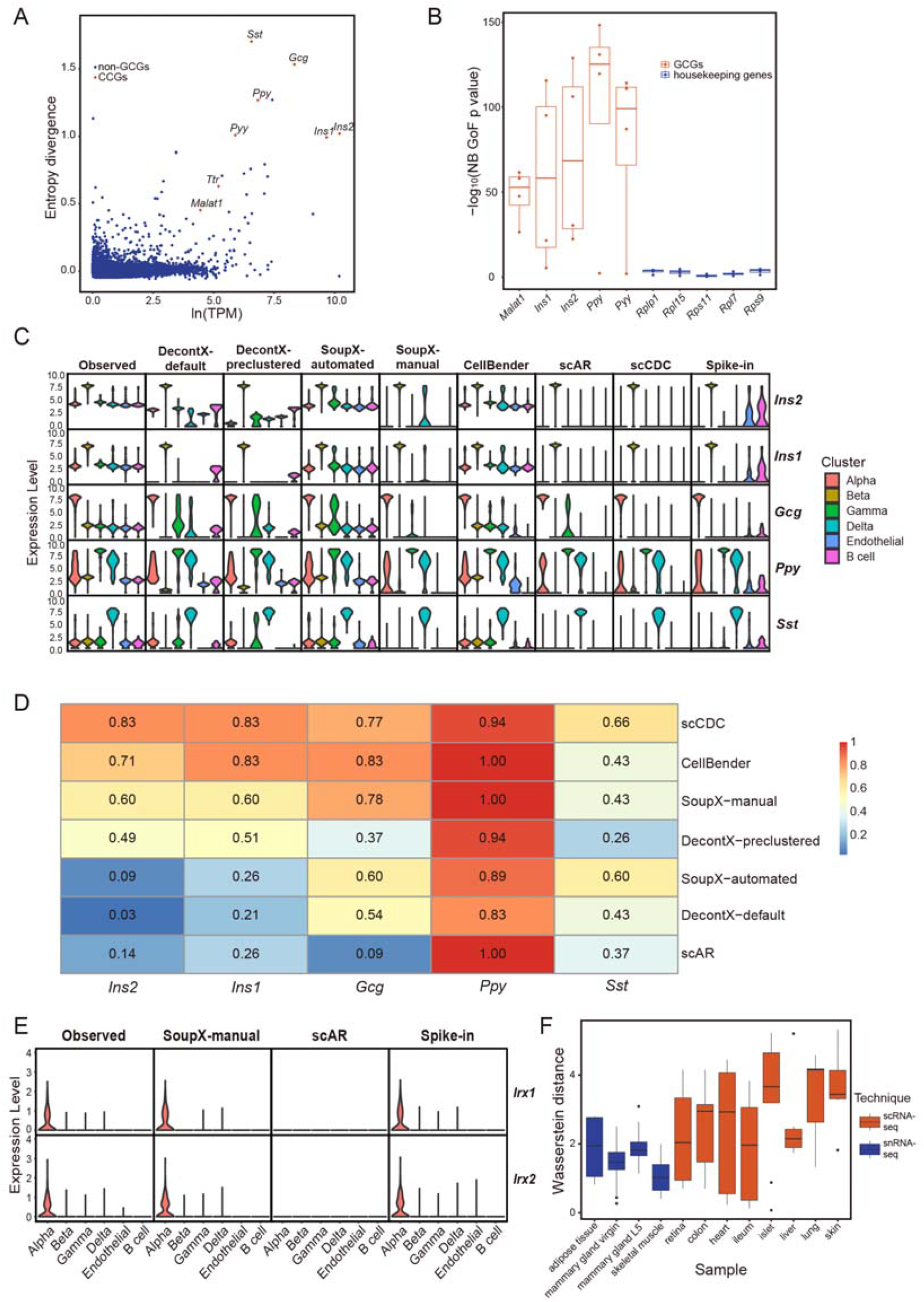
Contamination detection in a single-cell RNA-Seq dataset of mouse islet. (A) Scatter plot shows the relationship between the mean entropy divergence and the TPM (x-axis, log scale) of all the genes. GCGs are highlighted in red. (B) Box plot of the *p*-values of NB distribution goodness-of-fit test of GCGs and housekeeping genes. (C) Benchmarking scCDC, DecontX, SoupX, CellBender, and scAR in the single-cell RNA-Seq data in mouse pancreas islet with spike-in strategies. The violin plots show the normalized expression levels of the indicated GCGs before and after correction using the indicated methods by the default Seurat (V3). (D) The heatmap shows *Spearman*’s correlation between the mean expression levels of GCGs in the cell clusters in pancreas data after correction by different computational methods and by spike-in approach. (E) Feature loss in scAR. The violin plots show the normalized expression levels of two markers of alpha cells in pancreas, *Irx1* and *Irx2*, before and after correction using SoupX-manual, scAR, and spike-in approach by the default Seurat (V3). (F) Boxplot of the *Wasserstein* distances between the distribution of GCGs in the cell clusters that expressed the highest level and the estimated contamination distribution of GCGs in the indicated datasets. Mitochondria genes and erythroid cell-associated genes were removed in this analysis.

Previously, Sanchez et al. estimated the contamination fractions by including spike-in cells in the experiment and corrected the data based on the estimated contamination fractions [6]. We leveraged Sanchez et al.’s spike-in data to benchmark the correction performance of scCDC against that of DecontX, SoupX, CellBender, and scAR. Similar to their results on our in-house mammary gland snRNA-seq datasets, DecontX, SoupX-automated, and CellBender failed to correct the contamination. In contrast, scCDC, SoupX-manual, and scAR removed the contaminative counts of GCGs, resulting in even cleaner decontamination results than the spike-in-based correction, which did not remove all contaminative counts in the endothelial and B cell clusters (Figure 5C and Supplementary Figure 5). Notably, among all these computational methods, scCDC achieved the overall best correlations between its corrected counts and the spike-in-based corrected counts (Figure 5D).

Moreover, we checked if SoupX-manual and scAR over-corrected non-GCGs in the pancreas scRNA-seq dataset. Similar to our observations in our in-house mammary gland snRNA-seq data, SoupX-manual and scAR undesirably removed the counts of many housekeeping genes in more than 95% of cells (Supplementary Figure 6). In addition, we found that the reads of *Irx1* and *Irx2*, the marker genes of Alpha cells, were removed by scAR (Figure 5E), consistent with a previous report about potential gene loss caused by scAR [10].

**Figure 6.**
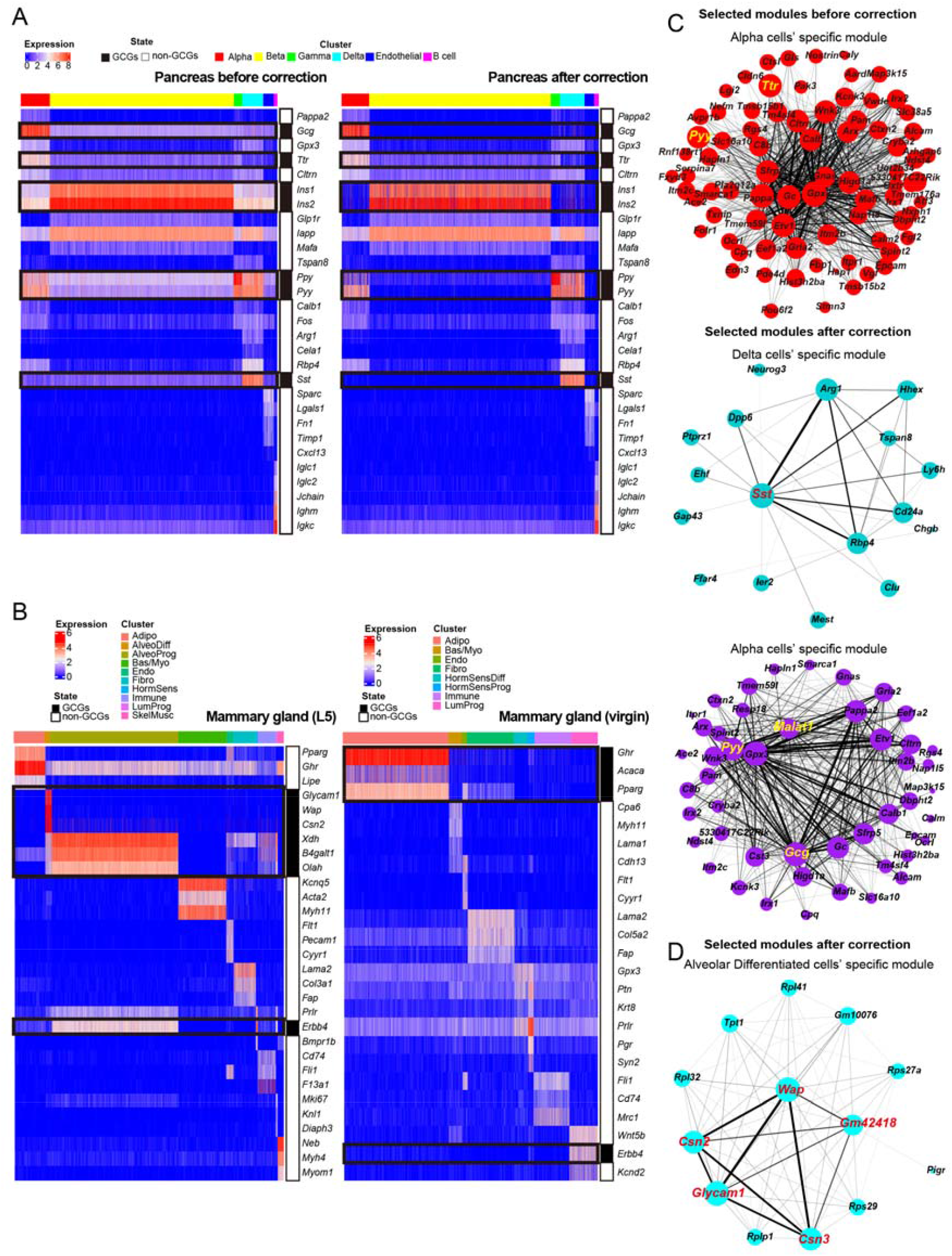
scCDC improves marker gene profiling and gene network analysis in mouse pancreas and mammary gland datasets. (A) Expression of selected top markers in each cell cluster is shown in the heatmaps after correction in mammary gland L5 (left) and mammary gland virgin (right) data. GCGs are highlighted on the right. (B) Comparison of marker gene expression before (left) and after (right) correction in pancreas dataset shown in heatmap. (C-D) The significant gene network modules associated with GCGs before and after correction were identified in pancreas (C) and mammary gland (D) data, respectively. The *Ttr*- and *Pyy*-centered module is derived from the uncorrected data, and the rest of the modules are derived from the corrected data.GCGs are highlighted.

Next, we applied scCDC to nine scRNA-seq and snRNA-seq datasets from various organs (Table 2) [6, 21-29]. Consistent with the results from the pancreas and mammary gland data, the identified GCGs are generally abundant in the organs and have pervasive contamination across cell types. Of note, the GCGs are mostly cell-type marker genes instead of housekeeping genes. We also noticed that hemoglobin genes, like *Hbas* and *Hbbs*, caused global contaminations in the adipose and heart tissues (Table 2), probably due to the lysis of erythroid cells, a common experimental step before preparing single-cell droplets. Our results suggest careful handling is needed when including this lysis step in scRNA-seq assays. Unlike a single-cell assay, a single-nuclei assay involves the procedure of cell nuclei extraction, which causes cellular RNAs to release and possibly leads to global contamination. Therefore, we speculated that contamination is more severe in snRNA-seq data than in scRNA-seq data. To test this hypothesis, we calculated the *Wasserstein* distance of each GCG’s count distributions in two groups of cells: eGCG-cells (where the GCG is expected to be unexpressed) and eGCG+ cells (where the GCG is expected to be expressed). Consistent with our hypothesis, the *Wasserstein* distances were overall smaller in the snRNA-seq datasets than in the scRNA-seq datasets (Figure 5F), confirming the more pervasive contamination in snRNA-seq data.

### scCDC’s contamination correction improves the accuracy of downstream analysis

Next, we examined the improvement of data analysis results after scCDC’s contamination correction. First, scCDC unmasked the expression patterns of cell-type marker genes, thus facilitating the identification of cell types. In the pancreas scRNA-seq dataset, scCDC revealed the exclusive expression of *Ins1* and *Ins2* in beta cells and *Gcg* in alpha cells (Figure 6A & Supplementary Figure 7A). In the lactating mammary gland (L5) snRNA-seq dataset, scCDC revealed the unique expression of milk protein genes, like *Wap* and *Csn2*, in AlveoDiff (differentiated alveolar) cells (Figure 6B and Supplementary Figure 7B). In the virgin mammary gland snRNA-seq dataset, scCDC showed the exclusive expression of adipocyte markers *Ghr, Acaca*, and *Pparg* and that of luminal progenitor marker *Erbb4* (Figure 6B and Supplementary Figure 7C).

Second, scCDC improved the construction of gene co-expression networks. We applied single-cell weighted gene co-expression network analysis (scWGCNA) [30] to the pancreas scRNA-seq dataset and the lactating mammary gland snRNA-seq dataset before and after scCDC’s correction (Supplementary Table 1). In the pancreas scRNA-seq dataset, many cell-type marker genes found as GCGs (such as delta-cell marker *Sst* and alpha-cell marker *Gcg*) were not identified in network modules before scCDC’s correction, suggesting that their contamination hindered the co-expression analysis. Only after scCDC’s correction, *Sst* and *Gcg* were identified as central genes of delta cells’ and alpha cells’ network modules, respectively (Figure 6C, Supplementary Figure 8A, and Table 1). In the snRNA-seq dataset of lactating mammary glands, the GCGs *Csn2, Csn3, Wap*, and *Glycam1* are well-known lactation-specific genes regulated by the same transcriptional machinery [31], but they were not identified in any network module before scCDC’s correction. In contrast, these four lactation-specific genes were identified as a network module of AlveoDiff cells after scCDC’s correction (Figure 6D, Supplementary Figure 8B, and Table 1). The results indicated that scCDC’s decontamination helped scWGCNA identify gene co-expression modules masked by the contamination.

## Discussion

Here, we developed a computational method, scCDC, to identify GCGs and correct the counts of GCGs independent of the availability of experimental spike-in controls or empty droplets. Our results indicate that global contamination warrants attention, especially for snRNA-seq assays, and scCDC effectively identified GCGs and corrected their contamination in scRNA-seq and snRNA-seq data. Compared to the existing computational methods, scCDC avoids the under-correction issue of DecontX, CellBender, and SoupX-automated and the over-correction issue of SoupX-manual and scAR, via the detection of GCGs (Table 1).

Among the existing computational methods, SoupX, CellBender, and scAR estimated the contaminative count distribution from empty droplets. However, these three methods have two limitations. First, it is too simplistic to assume that ambient RNA levels have the same distribution in empty droplets and in cell- or nucleus-containing droplets. The two reasons are (1) ambient RNAs are randomly distributed in empty droplets, but they may be attached to or absorbed by cells or nucleus in cell- or nucleus-containing droplets; (2) unlike cell- or nucleus-containing droplets, in empty droplets, the lack of endogenous RNAs may lead to more amplification of ambient RNAs and thus over-estimation of the contamination, e.g., the over-correction by SoupX and scAR on the pancreas scRNA-seq data (Figure 5E). Second, these methods are inapplicable to the processed gene-by-cell count matrices, which are common in public datasets and do not contain empty-droplet data.

In contrast, scCDC avoids these limitations by estimating the distribution of contaminated counts from real cells or nuclei, so scCDC can be applied to processed count matrices. Although DecontX can also be applied to processed count matrices, the performance of DecontX was worse than the other methods’ in our benchmark. We speculate that the DecontX algorithm’s convergence and iteration setting requires further optimization.

Note that scCDC and DecountX require the pre-clustering of cells, an issue we discussed in the Method Appendix. In contrast, SoupX, CellBender, and scAR do not require cell pre-clustering and should thus be more suitable for correcting the data composed of obscure cell populations (Table 1).

What distinguishes scCDC from the existing methods is that scCDC detects GCGs and only corrects the expression counts of GCGs. This strategy, which was also used in scImpute for the imputation problem, minimizes data alteration to avoid the over-correction issue of SoupX and scAR [32]. This strategy makes scCDC conservative so that cell clusters only change slightly after scCDC’s correction. Note that the GCG detection algorithm in scCDC may be further optimized by using alternative strategies for estimating the expected entropy-expression curve in each cell cluster. We will consider relevant strategies in our maintenance of the scCDC software package.

Similar to scRNA-seq and snRNA-seq, single-cell proteomics was also found to have contamination [11]. According, decontamination methods such as dbs was developed [33]. Although we focused on correcting the contamination in scRNA-seq and snRNA-seq datasets in this study, scCDC is also applicable to single-cell proteomics data theoretically. The performance of scCDC on single-cell proteomics data can be benchmarked in a future study.

## Conclusions

Global contamination by ambient RNAs is ubiquitous in single-cell and single-nuclei RNA-seq assays. We proposed scCDC as a computational method to detect global contamination-causing genes and corrects these genes’ expression data. The gene-specific correction strategy makes scCDC more sensitive to global contamination and less likely to over-correct housekeeping genes, compared to the existing computational methods. Data correction by scCDC improves marker gene identification and gene network construction.

## Methods

### Calculation of cell-cluster-specific gene entropy divergences in scCDC

For gene *g* in cell cluster *c*, the **entropy** is defined as 

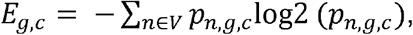

where *V* is the set of unique values in *v*_*g,c*_′, a vector of gene *g*’s counts in the cells in cluster *c* ; *p*_*n,g,c*_ is the frequency of the count value *n* in *v*_*g,c*_′ defined as 

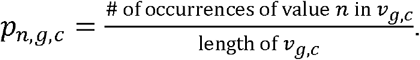

For cell cluster *c*, the following procedure is used to calculate the cell-cluster-specific **expected entropy-expression curve**, inspired by the ROGUE score in [34].

1. Calculate each gene *g*’s mean expression in cell cluster *c* as

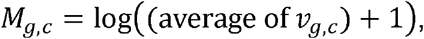

and calculate *E*_*g,c*_, defined above.
2. For *b*=1,…,10, in the *b*-th subsampling run, do the following.
  i. Randomly sample 80% of genes.
  ii. Use the R function smooth.spline() to fit a curve between the sampled genes’ entropy values (y; response variable) and mean expression values (x; explanatory variable), using the following R code:

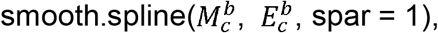

where 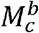 is a vector containing the randomly sampled ge nes’ *M*_*g,c*_ values, and 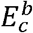 is a vector containing the randomly sampled genes’ *E*_*g,c*_ values. Denote the fitted curve by function 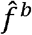 that maps a gene’s mean expression to entropy.
  iii. For each sampled gene *g*, calculate the residual 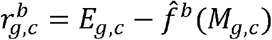, i.e., the difference between the gene’s entropy and the fitted entropy from step ii. Pool all residuals into a vector 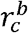. Assuming all residuals follow a normal distribution, define gene *g* as an outlier if its residual falls into the top 1% tail of the fitted normal distribution, i.e., using R code, if 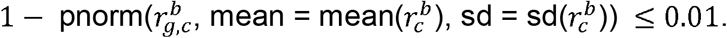
  iv. Remove the outlier genes detected in step iii and refit the curve as in step ii.
  v. Detect outlier genes as in step iii based on the refitted curve in step iv.
  vi. Remove the outlier genes detected in step v and refit the curve as in step ii.
  vii. Output the curve from step vi.
3. Calculate the expected entropy-expression curve by averaging the 10 curves from the subsampling runs. Specifically, for each gene *g*, its expected entropy is the average of the 10 fitted entropy values.

Finally, the **entropy divergence** of *g* in cell cluster *c* is defined as 

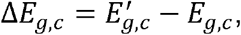

where 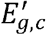 is the expected entropy of gene *g* in cell cluster *c*, calculated based on gene *g*’s average expression and the expected entropy-expression curve in cell cluster *c*. Since we expect that gene *g*’s ambient RNAs would deflate its entropy *E*_*g,c*_, a large and positive Δ*E*_*g,c*_, would indicate severe contamination of gene *g* in cell cluster *c*.

### GCG Identification in scCDC

Small cell clusters (with fewer than 100 cells) are not considered in this GCG identification step. Figure 2B illustrate the GCG identification procedures described below.

1. Among the considered cell clusters, in every cluster *c*, the genes with “significantly” large entropy divergences (and have non-zero counts in at least 80% of cells in the cluster) would be identified as the **candidate GCGs** of cluster *c*. Specifically, we fit a normal distribution of all genes’ entropy divergences, denoted by the vector Δ*E*_c_. Then we calculate a pseudo-p-value of gene *g*, denoted by *pp*_*g*_, as

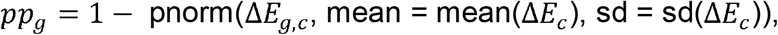

and set a 0.05 threshold on the adjusted pseudo-p-values based on the Benjamini-Hochberg procedure. That is, any gene *g* whose post-adjustment *pp*_*g*_ ≤ 0.05 would be called a candidate GCG in cluster *c*, if gene *g* is expressed in at least 80% of the cells in cluster *c*.

2. Across the considered cell clusters, the genes found as candidate GCGs in at least 80% (referred to as the **restriction factor**, which can be user-specified; the selection of an appropriate restriction factor is discussed in the Method Appendix) of the clusters would be found as the **GCGs**. In other words, the GCGs are the genes that are stably found as candidate GCGs in many clusters.

### Estimation of a GCG’s contaminative count distribution in scCDC

Each GCG’s contaminative count distribution is estimated by the GCG’s counts in the cells that are not expected to express the GCG endogenously (i.e., **eGCG-cells**; illustrated in Figure 2B). A GCG’s eGCG-cells are defined by pooling the cell cluster with the lowest median expression of the GCG (i.e., the most eGCG-cluster) and the other cell clusters whose 1-D Wasserstein distances to the most eGCG-cluster are less than 1. Specifically, given the GCG, the 1-D Wasserstein distance between two cell clusters is calculated as

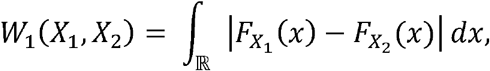

Where *X*_1_ and *X*_2_ are random variables denoting the GCG’s normalized and log-transformed counts (adopted from Seurat) in the two clusters, 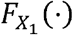 and 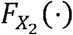 are the cumulative distribution functions, and the calculation of W_1_(*X*_1_,*X*_2_) is done using the ‘transport’ (v0.12-2) package developed by Schuhmacher et al., 2019 (https://cran.r-project.org/package=transport). The 1-D Wasserstein distance threshold of 1 is chosen as the medium of the maximum 1-D Wasserstein distances of expressed (detected in more than 50% cells) housekeeping genes (from a list of 70 housekeeping genes) between all cluster pairs in six datasets (i.e., each included housekeeping gene has a maximum distance in each dataset; see Supplementary Figure 10). A GCG’s **eGCG+ cells** are expected to have the GCG endogenously expressed, and they are defined by excluding the eGCG-cells.

### Correction of a GCG’s contaminative counts by scCDC

Given a GCG, the following two approaches are used to correct the GCG’s counts, depending on the number of clusters containing the GCG’s eGCG+ cells.

1. **Otsu’s method-based correction** is applied when only one cell cluster contains eGCG+ cells. First, we balance the numbers of eGCG+ cells and eGCG-cells by oversampling. Specifically, denoting by *n* the maximum of the number of eGCG+ cells and the number of eGCG-cells, we oversample (i.e., sample with replacement) the smaller group of cells (e.g., eGCG-cells) so that both the eGCG+ and eGCG-groups have *n* cells. Pooling the GCG’s counts in the 2*n* cells, we apply the Otsu’s method to a threshold *t* such that the weighted sum of intra-class variance

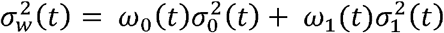

is minimized, where ω_0_ (*t*)is the proportion of the counts smaller than *t* (among the 2*n* counts), 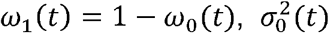 is the variance of the counts smaller than *t*, and 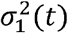 is the variance of the counts greater than or equal to *t*. Given the threshold *t*, we correct the GCG’s count in every cell by subtracting *t*, with a truncation at zero so that the GCG’s minimum count is zero.

**(2) Cumulative distribution function (CDF)-based correction** is applied when more than one cell cluster contains the GCG’s eGCG+ cells.

For any GCG *i* in cell *j* of cluster *c*, denoting by *x*_ij_ the observed count, we define the corrected count 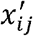 as

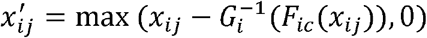

where *F*_*ic*_ is the CDF of GCG *i*’s counts in cell cluster *c*, and 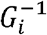 is the inverse CDF of GCG *i*’s counts in the eGCG-cells.

Together, scCDC’s tuning parameters are summarized in Table 3.

**Table 3.** Tuning parameters in the functions of scCDC.

| Function | Parameter | Description | Default |
| --- | --- | --- | --- |
| Contamination Detection | restrict_factor | The minimum proportion of cell clusters in which a GCG is found as a candidate GCG | 0.8 |
|  | min.cells | The minimum cell number of the cell clusters used for finding candidate GCGs | 100 |
| Contamination Correction | was.thres | The threshold of the 1-D Wasserstein distance used to define eGCG- clusters | 1 |

### Evaluation of Otsu’s method versus CDF-based method

We first simulated an array of uncontaminated counts with a mixture of two negative binomial distributions, one of which represents the eGCG+ cluster (size = 1, µ = 1000) while the other one represents the eGCG-cluster (size = 1, µ = 1). Then, we built negative binomial distributions as contamination distributions with various means that represent various contamination levels (size = 10, µ = 0 – 500). For each contamination level, we sampled values from the contamination distribution, which were randomly added to the uncontaminated counts to obtain the array of contaminative counts.

To compare Otsu’s method with CDF-based method, these two correction methods were applied to the array of contaminative counts after a pre-clustering process using the k-means algorithm (k = 2). *Spearman’s* correlations were calculated between corrected counts and uncontaminated counts for each method under all of the contamination levels mentioned.

### Count correction by SoupX, DecontX, CellBender, and scAR

SoupX (v1.5.2), DecontX in Celda (v1.10.0), CellBender (v0.3.0), and scAR (v0.4.3) were employed for count correction.

For SoupX, both raw feature matrix and filtered feature matrix generated by Cellranger (v6.0.1) were used to create the Soup Channel object, followed by standard correction workflow in the guidance [5]. The “automated” and “manual” approaches were applied, respectively. The identified GCGs were provided as the ‘non expressed genes’, whose RNAs in specified cells were treated as solely ambient ones, in the “manual” mode.

For DecontX, the correction was applied to the filtered feature matrix in the datasets. The default procedure, referred to as the “default” mode, was first performed.

Alternatively, the pre-clustering information obtained from Seurat [35] was provided manually in the “pre-clustered” mode.

For CellBender, the correction used the raw feature matrix with the remove-background function following the tutorial (https://cellbender.readthedocs.io/en/latest/getting_started/remove_background/index.html).

For scAR, both raw feature matrix and filtered feature matrix were used to perform the pipeline based on the tutorial (https://scar-tutorials.readthedocs.io/en/latest/tutorials/scAR_tutorial_mRNA_denoising.html). The filtering scale was applied to the filtered feature matrix of datasets as listed in Supplementary Table 2.

### Generation of simulated PBMC single-cell dataset

To generate the simulated PBMC single-cell dataset, we first obtained a real PBMC dataset’ pbmcsca.SeuratData’ from the SeuratData R package (https://github.com/satijalab/seurat-data). We then sub-selected the dataset generated by the 10x Chromium (v2) technology under the experiment’ pbmc2’, using the ‘meta. data’ information from ‘pbmcsca.SeuratData’. Next, we filtered out the ERCC spike-in’s, the mitochondrial genes, and the gene *MALAT1*, and we selected five cell types (B cells, CD14+ monocytes, natural killer cells, CD4+ T cells, and cytotoxic T cells). Using the filtered and sub-selected real dataset from above, we applied the simulator scDesign2 [19, 36] to fit one multivariate probabilistic model to each of the five cell types.

The resulting gene expression matrix was stored as the file sce_10x_pbmc2_hca_corrected.rds.

The sce_10x_pbmc2_hca_corrected.rds file and the code for reproducing it are available at https://zenodo.org/record/6395574#.YrXp5JPMKEt. In particular, the rds file is under Code summary.zip/Figure 7 and supplementary S3 to S10/imputation_comparison_0614/Data_gen/data/; the code is under Code summary.zip/Data simulation/. The rds file can be generated by sequentially executing the seven steps in the code directory.

### Artificial contamination of simulated PBMC single-cell dataset

To further simulate global contamination, an additional PBMC dataset was generated by randomly blending the original raw count matrix with an artificial contaminative count matrix composed of three marker genes of CD14+ monocytes, *S100A9, S100A8*, and *LYZ*. If not indicated, the contaminative count matrix was generated following negative binomial distributions using the mean of the average count of the original raw count matrix and a size of 10. Then a contaminative count was randomly selected from the matrix and added to each original raw count. Alternatively, contaminative counts were generated using a fixed mean and size of 1, 10, 50, 100.#x0026;

### Single-nuclei RNA-seq in mammary glands

Eight-week-old female C57BL/6N mice were timed mated. Abdominal and thoracic mammary tissues from nulliparous mice (virgin) and mice at lactation day 5 (L5) were harvested and lymph nodes in abdominal mammary tissues were removed. Mammary tissues were snap-frozen in liquid nitrogen followed by nuclei extraction and single-nuclei RNA sequencing (snRNA-seq) on a 10X Genomics platform in Lianchuan Biology Technology Co.

### General single-cell and single-nuclei data processing

Cellranger (v6.0.1) was used to map raw reads to mouse or human reference genomes and obtain raw and filtered count matrixes of genes. Seurat (v4.0.3) was used for data filtration, principal component analysis (PCA), dimension reduction, clustering, marker gene identification and data visualization. Specifically, for each dataset, cells with insufficient genes, molecules and high mitochondria gene percentage were first filtered. The data were then normalized and top variable genes were identified. Scaling, dimension reduction, and clustering were then performed. The specific parameters for filtering, dimension reduction, and clustering used in each dataset are provided in Supplementary Table 2.

When benchmarking for pre-clustering, the top 1000 variable genes identified by SeuratVST [35], scPNMF (v1.0) [37], and Scater (v1.20.1) [38] were used for dimension reduction, respectively. Rand index (ARI) values were calculated as described previously [37].

Visualization of clusters and gene expression and marker gene identification was done in Seurat. Weighted gene co-expression network analysis (WGCNA) was done using the scWGCNA (v1.0.0) package [30].

## Declarations

The authors declare that they have no competing interests.

### Ethics approval and consent to participate

All animal experiments were carried out according to the Institutional Ethical Guidelines on animal care and were approved by the Ethics Committee of The Second Affiliated Hospital Zhejiang University School of Medicine (2019-421).

## Availability of data and materials

The scCDC R package is available at https://github.com/ChaochenWang/scCDC. The processed datasets and the code scripts used to generate the figures are available on Zenodo (DOI:10.5281/zenodo.6905189).

The published single-cell and single-nuclei datasets were downloaded from the GEO database and Human Cell Atlas with the accession numbers listed in Table 2. The single-nuclei RNA-seq data of mouse mammary glands are deposited in the Genome Sequence Archive in National Genomics Data Center, China National Center for Bioinformation / Beijing Institute of Genomics, Chinese Academy of Sciences (GSA: CRA007450).

The total RNA-seq data in mouse lactating and virgin mammary glands are from the GEO database with accession numbers: GSE115370 [39] and GSE52016 [40], respectively. The total RNA-seq data from mouse pancreatic islets were downloaded from GEO: GSE148809 [41]. The genes encoding secrecting proteins were predicted in SignalP 4.0 [42], and the list of protein-coding genes were obtained from the Refseq database [43].

## Authors’ contributions

W.C. conceived and designed the project. W.W., C. Y. and L. Z developed the scCDC pipeline and performed data analysis. W.C., W.W., C. Y. and L. Z wrote the manuscript. W.C. and L. J. J. supervised the project and edited the manuscript. Xu Y. conducted the snRNA-seq experiment in the mammary gland and analyzed the data. S.T. generated the simulated dataset. Xiao Y. and L. W. edited the manuscript.

## Acknowledgments

The authors would like to thank the suggestions from Prof. Kuan Yoow Chan and the support from the animal facility of ZJU-UoE Institute of Zhejiang University.

