## Supplementary material for "scCDC: a computational method for gene-specific contamination detection and correction in single-cell and single-nucleus RNA-seq data": Method Appendix

**Sensitivity analysis of scCDC to the resolution of cell pre-clustering**

Pre-clustering is a crucial step for the performance of DecontX [1]. Similar to DecontX, scCDC requires a pre-clustering step. To investigate the impacts of pre-clustering resolution on scCDC’s performance, we clustered the cells in the pancreas scRNA-seq dataset with different resolutions by the Louvain algorithm in Seurat V3 [2]. When the number of clusters increased from 5 to 10 (Louvain resolution from 0.1 to 0.4), scCDC identified only one more GCG (Supplementary Figure 9A)

In the virgin mammary gland snRNA-seq dataset, increasing the Louvain resolution from 0.1 to 0.4 decreased the number of identified GCGs from 38 to 10, with 9 GCGs in common (Supplementary Figure 9B). The reason is that more clusters made fewer genes satisfy the default 80% restriction factor (i.e., GCGs must have statistically significant entropy divergences in more than 80% of the cell clusters; details in Methods). Hence, when the restriction factor was reduced to 60% at the Louvain resolution 0.4, more GCGs were detected (Supplementary Figure 9B). Notably, the GCGs identified in the settings were significantly overlapped (Supplementary Figure 9C). Taken together, these results indicated that the GCG identification by scCDC is sensitive to the pre-clustering resolution and an optimal restriction factor should be selected given a pre-clustering resolution.

**scPNMF assists scCDC in selecting the restriction factor given the pre-clustering resolution**

Therefore, we pursued to find a method to assist selecting the restriction factor in scCDC when a pre-clustering is performed with a given resolution. Because single-cell analysis largely relies on cluster-informative genes as shown in Figure 6, the targets of contamination correction should focus on these genes. Therefore, we reasoned that an appropriate pre-clustering setting is ought to maximize the proportion of cluster-informative genes in GCGs. However, in the contaminated pancreas and virgin mammary gland datasets, the widely used algorithms Seurat and Scater failed to identify GCGs as top informative genes, although most of which were known marker genes of various cell types. Recently, we developed scPNMF to unsupervisedly select cluster-informative genes in scRNA-seq data [3]. Unlike Seurat and Scater, scPNMF successfully identified the GCGs as the top 10 informative genes (Supplementary Figure 9D). Next, we examined the proportion of top 200 cluster-informative genes in the GCGs of the virgin mammary gland dataset, when different resolution and restriction factor were set in pre-clustering. The result showed that the combinations of resolution/restriction factor of 0.1/0.8 and 0.4/0.6 led to maximal proportion of cluster-informative genes in identified GCGs (Supplementary Figure 9E). As expected, most of the GCGs were overlapped (Supplementary Figure 9F). These data indicated that scPNMF is a robust method in cluster-informative gene identification in contaminated datasets. It is suggested that scPNMF-identified top 200 cluster-informative genes an assist scCDC in selecting the restriction factor given a pre-clustering resolution.
