## Supplementary Figures for "scCDC: a computational method for gene-specific contamination detection and correction in single-cell and single-nucleus RNA-seq data"

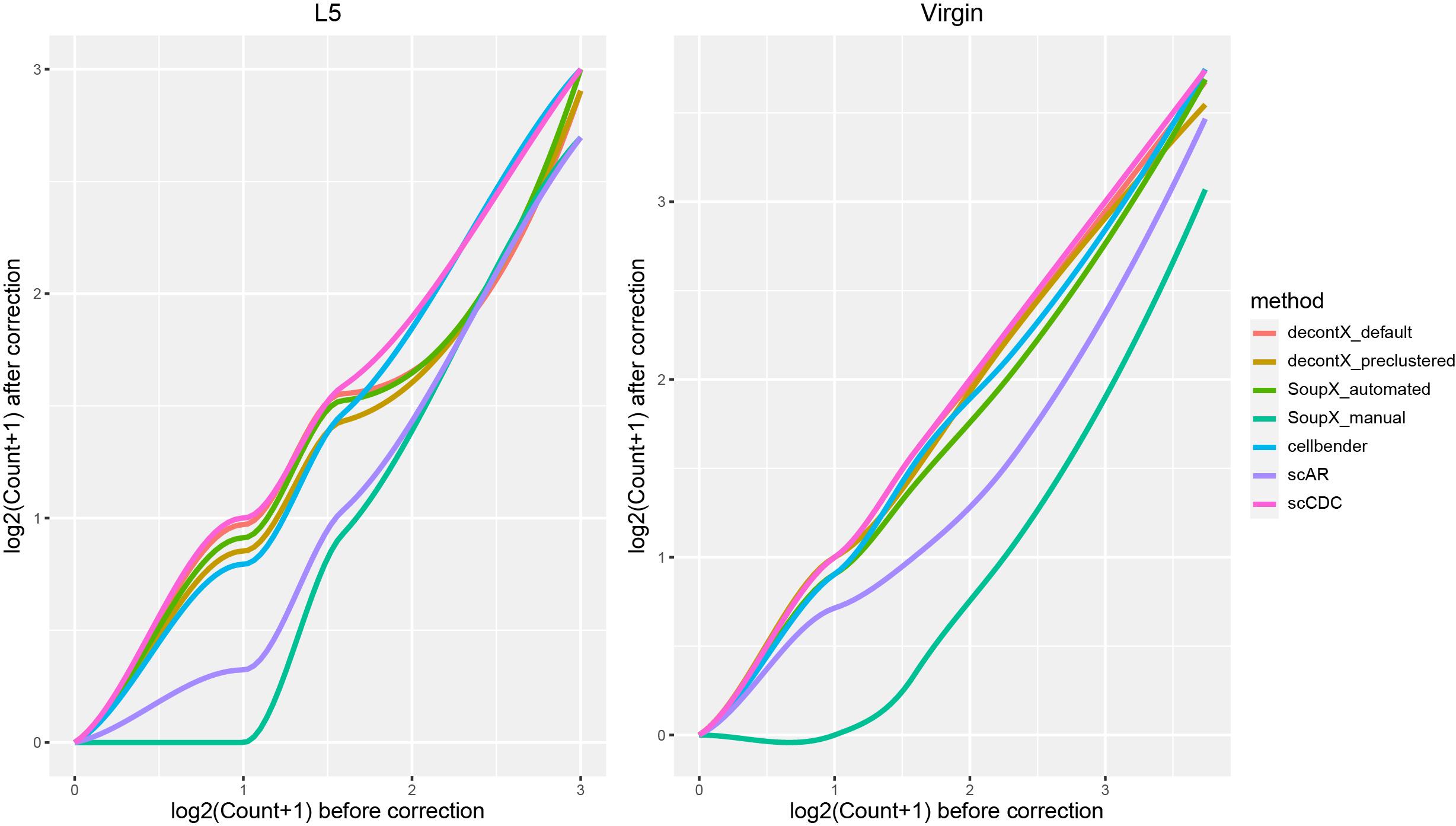
**Supplementary Figure 1.** Overcorrection by SoupX and scAR in snRNA-seq mammary gland datasets. Q-Q plots that compare the distribution of 66 housekeeping genes’ 95^th^ percentile counts in all cells (one percentile per gene) after correction by each method vs. the distribution before correction in lactating (left) and virgin (right) mammary gland snRNA-seq datasets.

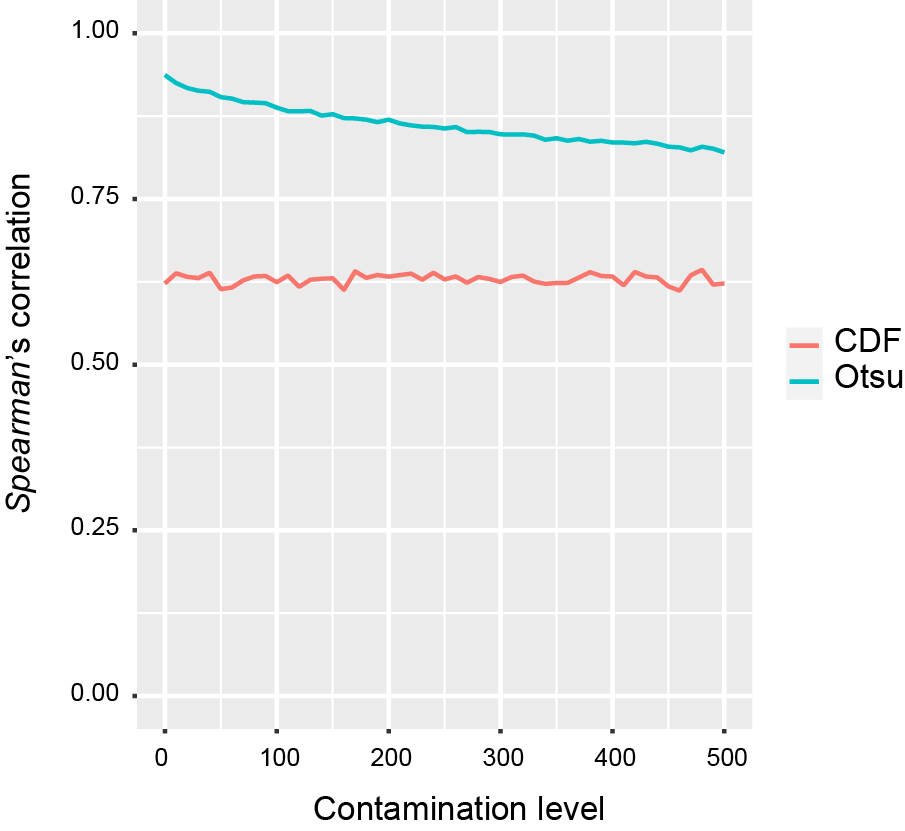

**Supplementary Figure 2.** *Otsu*’s method outperformed CDF-based method for correcting simulated contaminative counts in a mixture of one eGCG+ and one eGCG- cell cluster. The empirical curve shows the *Spearman*’s correlation between corrected counts and uncontaminated counts relative to the contamination level as in Figure 3B.

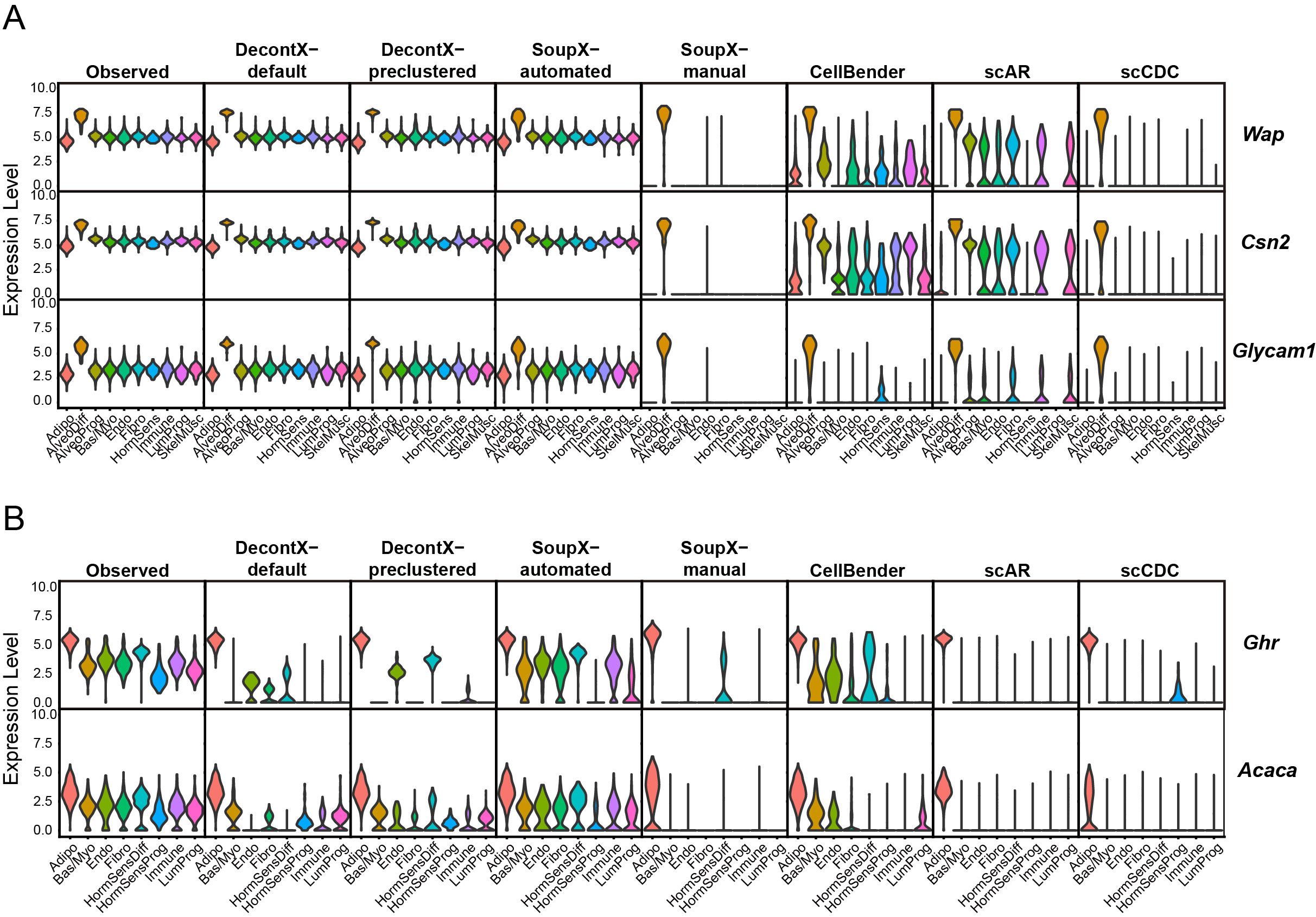
**Supplementary Figure 3.** Benchmarking scCDC, DecontX, SoupX, CellBender, and scAR in the snRNA-Seq datasets of mammary glands (virgin and L5). The violin plots show the expression levels of the indicated GCGs in L5 mammary gland (A) and virgin mammary gland (B) before and after correction using the indicated methods.

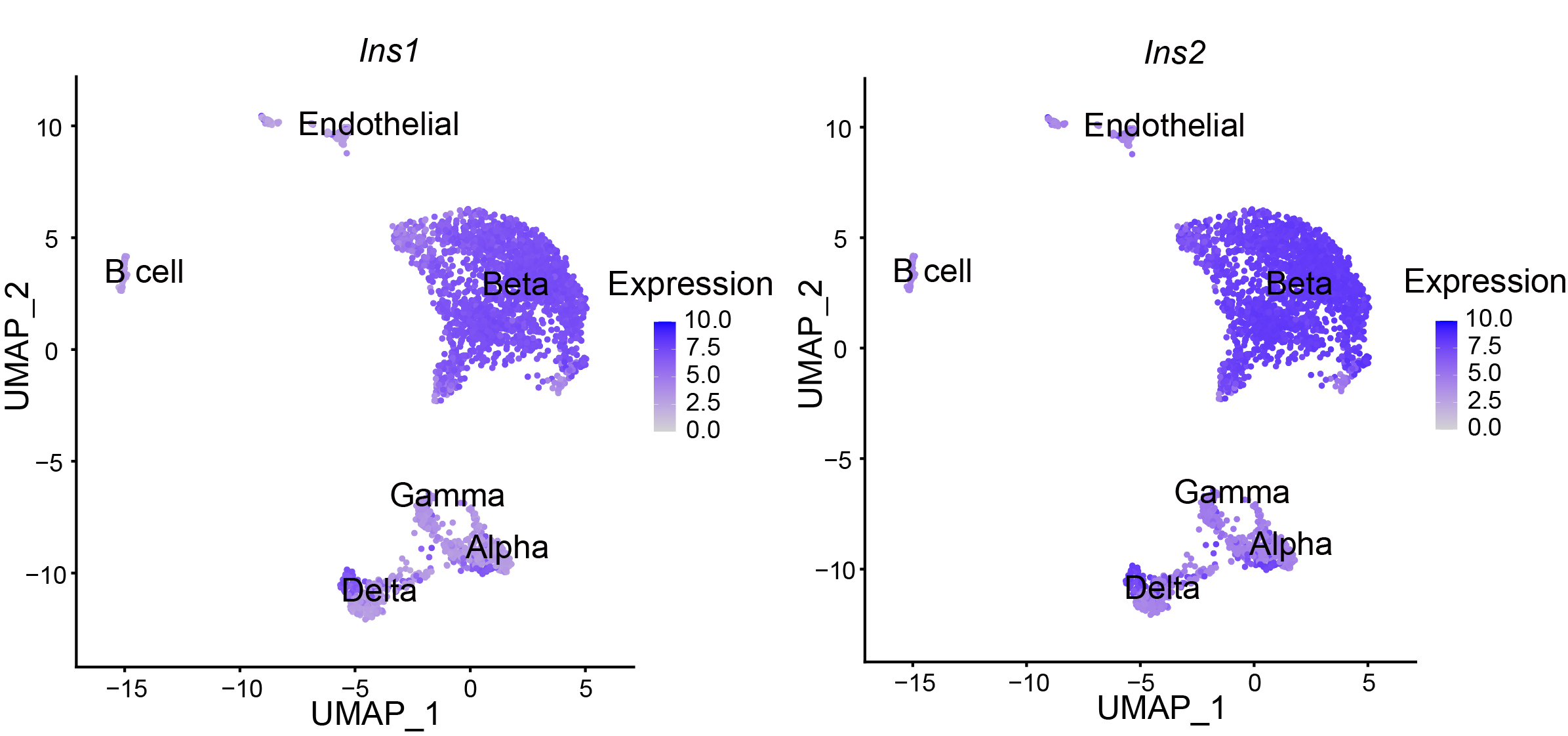

**Supplementary Figure 4.** Contamination in a scRNA-Seq dataset of mouse islet. The expression of beta cell markers *Ins1* and *Ins2* in the cells are shown in the UMAP plots.

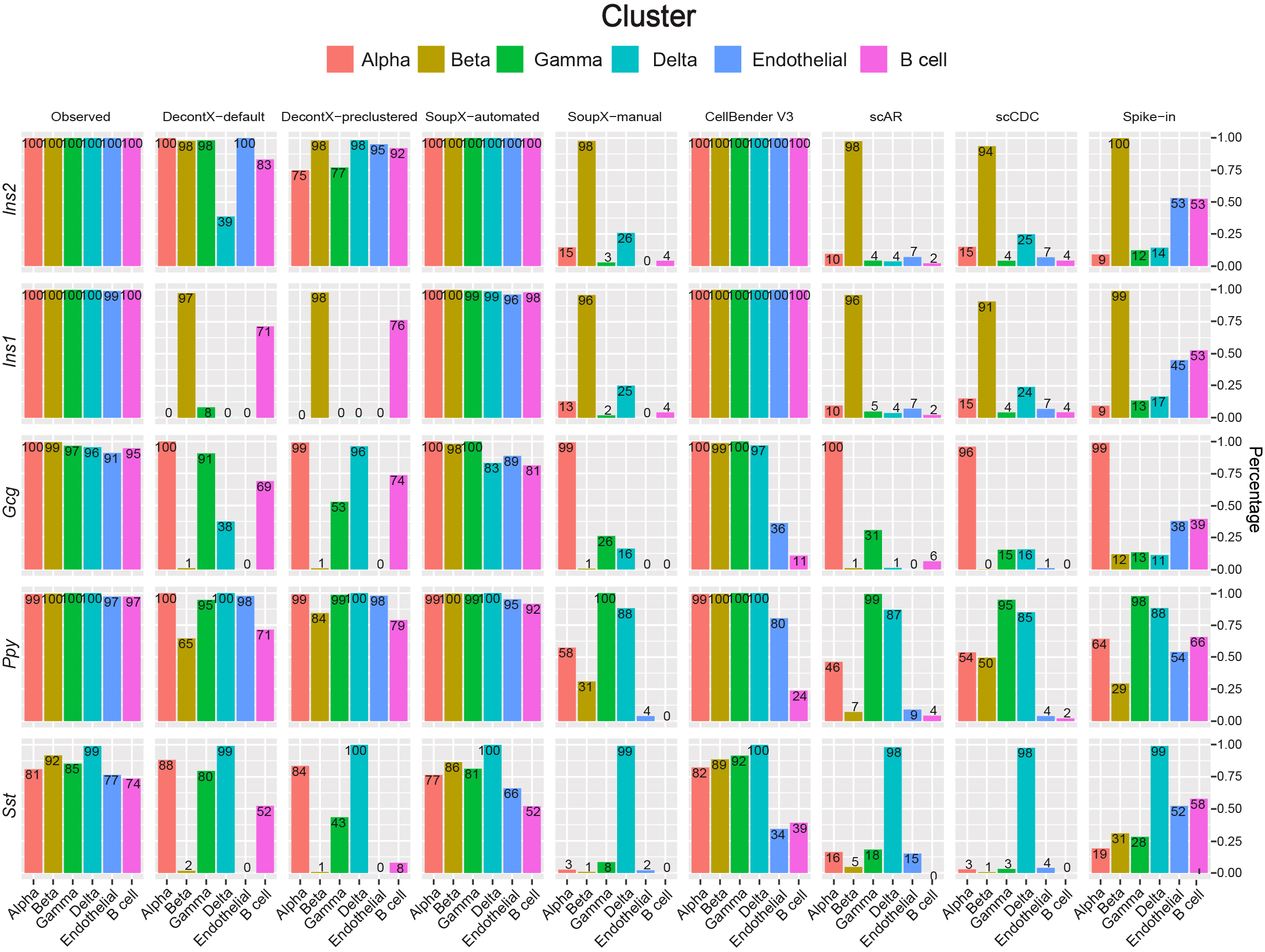

**Supplementary Figure 5.** Benchmarking scCDC, DecontX, SoupX, CellBender, and scAR in the scRNA-Seq data of mouse pancreas islet with spike-ins. The barplots show the percentages of cells expressing GCGs in the cell clusters before and after correction using the indicated methods.

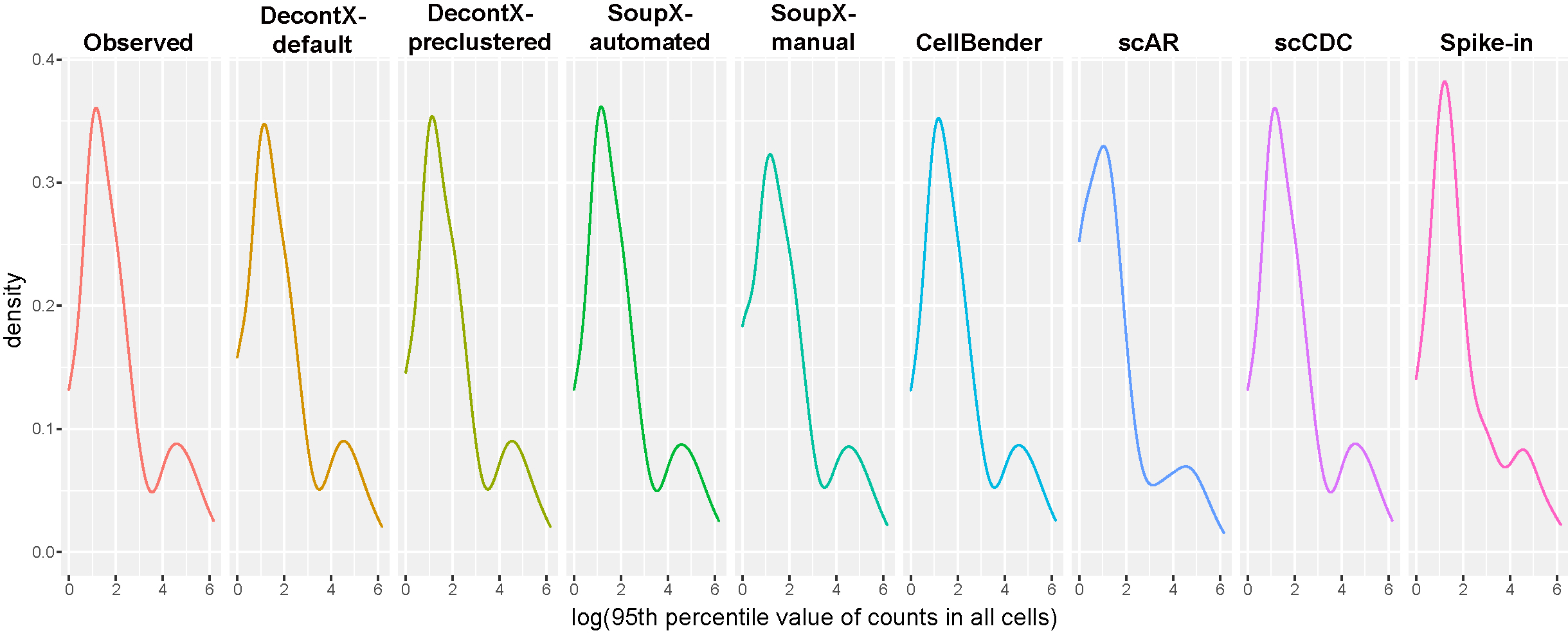

**Supplementary Figure 6.** Overcorrection by SoupX and scAR in pancreas scRNA-seq dataset. Density plots of the 95^th^ percentile value of counts of 66 housekeeping genes in all the cells before and after correction by the indicated methods in the pancreas scRNA-seq datasets.

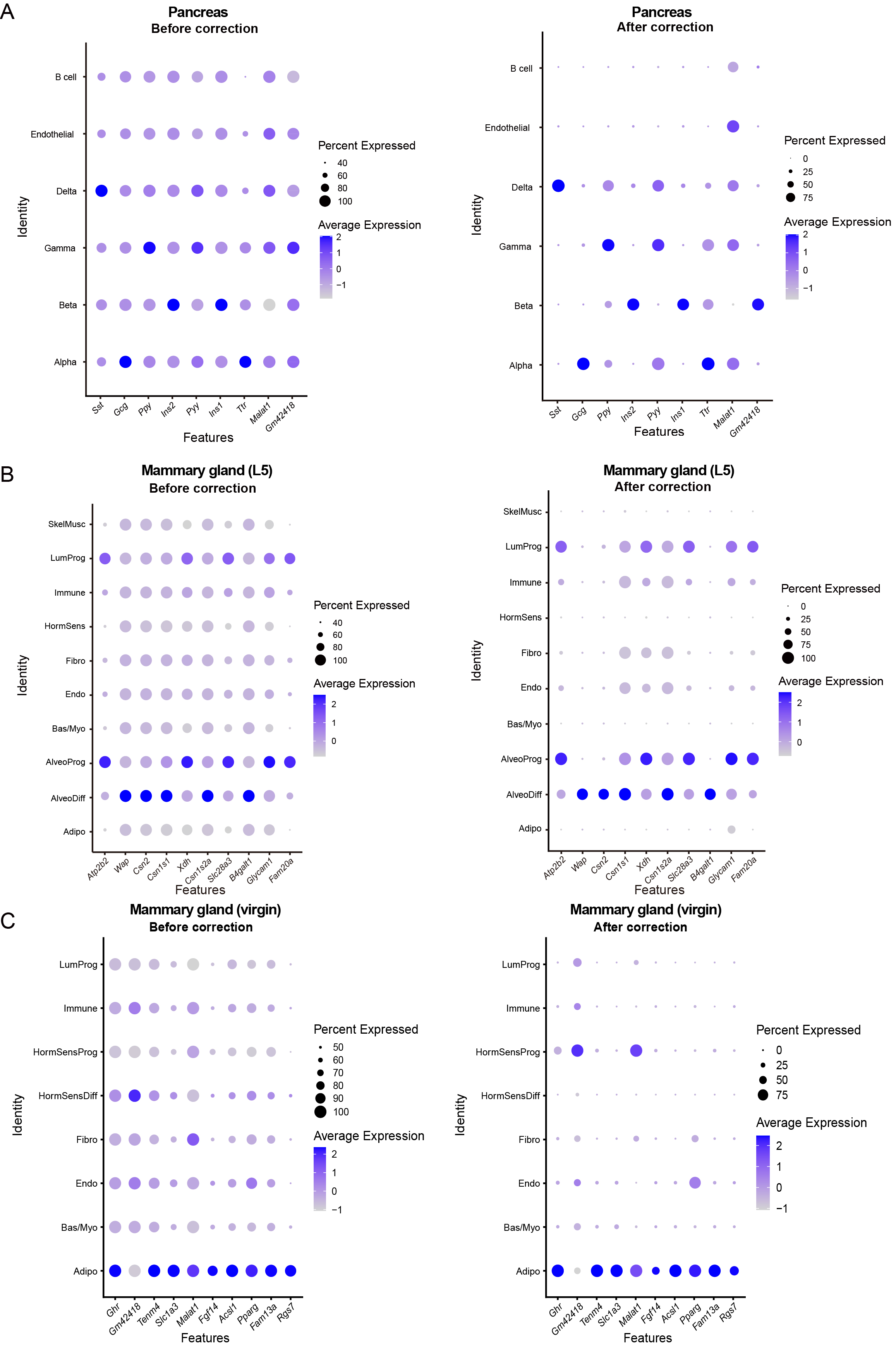
**Supplementary Figure 7.** scCDC corrected GCGs’ expression in mouse pancrea and mammary gland data. Dotplots show the expression of selected GCGs in pancreas (A), L5 mammary gland (B) and virgin mammary gland (C) before (left) and after correction (right), respectively.

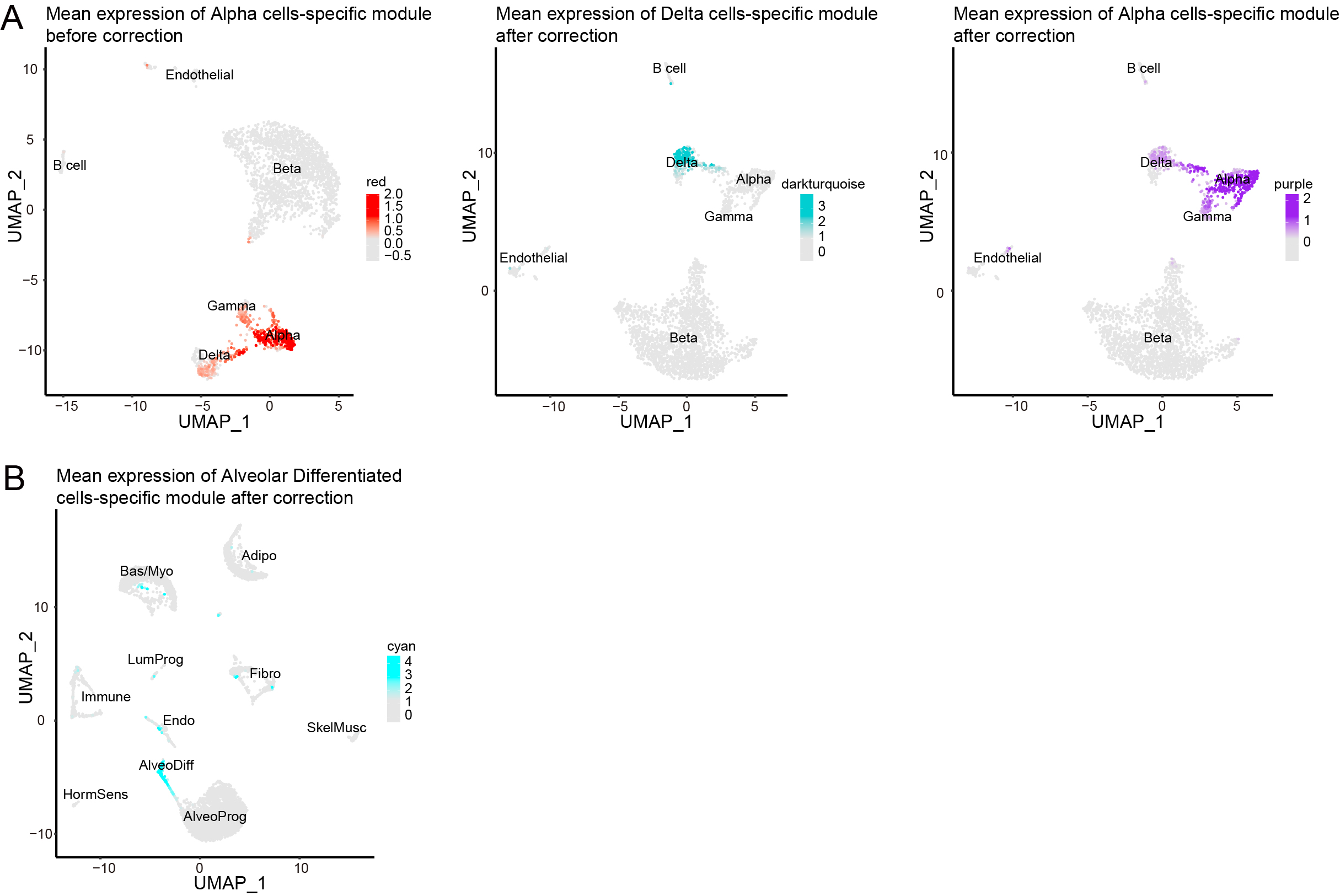
**Supplementary Figure 8.** The distribution of module signals was consistent with gene identities and the clustering of cells was hardly altered

(A-B) Distribution of mean expression of module genes in cell clusters in the pancreas (A) and mammary gland data (B) before and after correction in Figure 5.

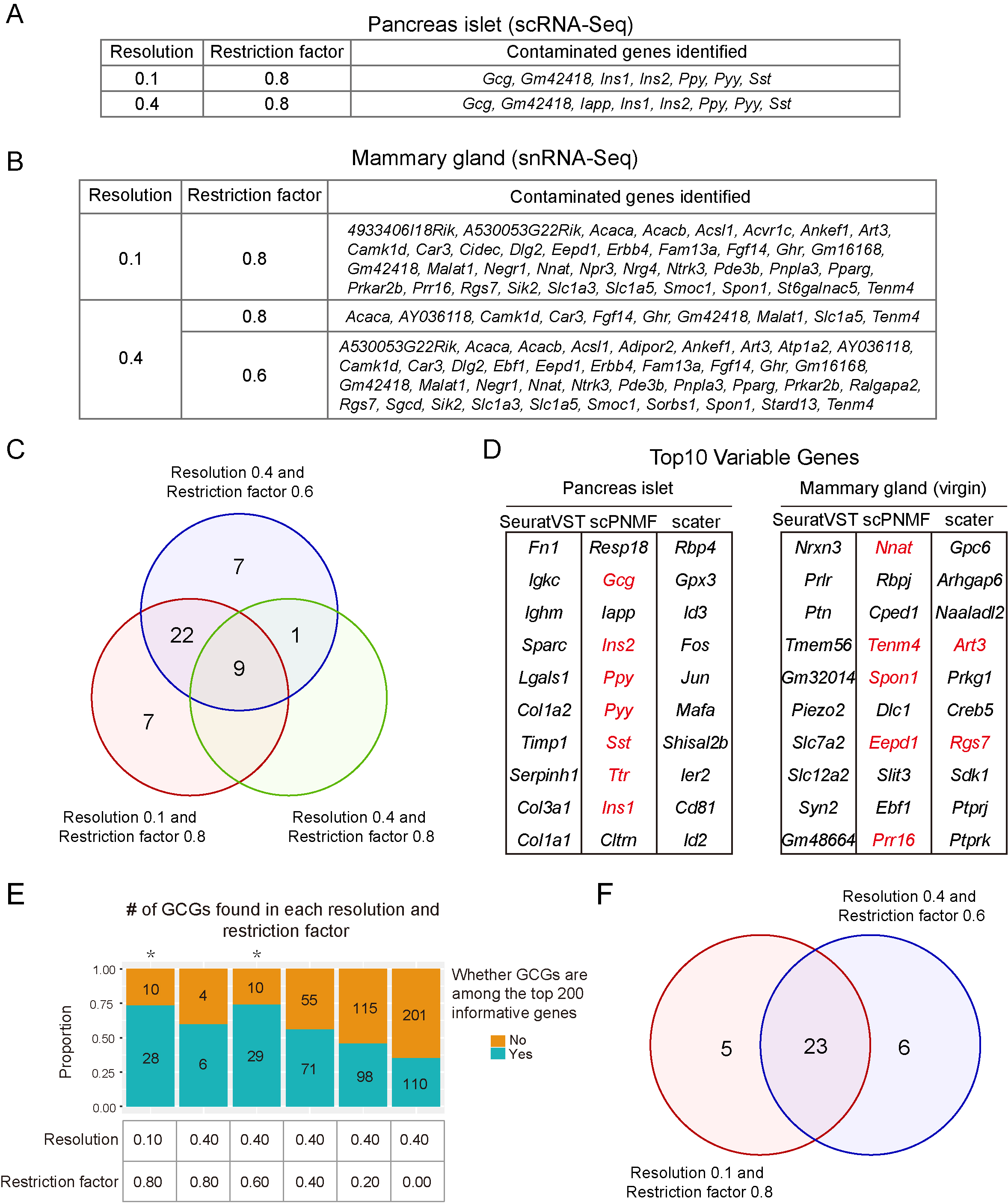

**Supplementary Figure 9.** scPNMF assists pre-clustering setting and GCG identification in contaminated datasets. (A) The list of GCGs identified by scCDC after pre-clustering with different resolutions in the mouse pancreas islet data. (B) The list of GCGs identified by scCDC after pre-clustering with different resolutions and restriction factors in the virgin mouse mammary gland snRNA-Seq data. (C) Venn diagram shows the intersection of GCGs that are detected under different resolutions and restriction factors. (D) Top 10 variable genes identified by SeuratVST, scPNMF, and Scater methods in pancreas islet scRNA-Seq and virgin mammary gland snRNA-Seq data. GCGs defined by scCDC are highlighted. (E) The barplot shows the percentage of scPNMF-determined cluster-informative genes in GCGs identified with indicated settings in the virgin mammary gland dataset as in (B). (F) Venn diagram shows the intersection of GCGs that are simultaneously identified as top 200 informative genes by scPNMF. Different groups of GCGs are detected under different resolutions and restriction factors.

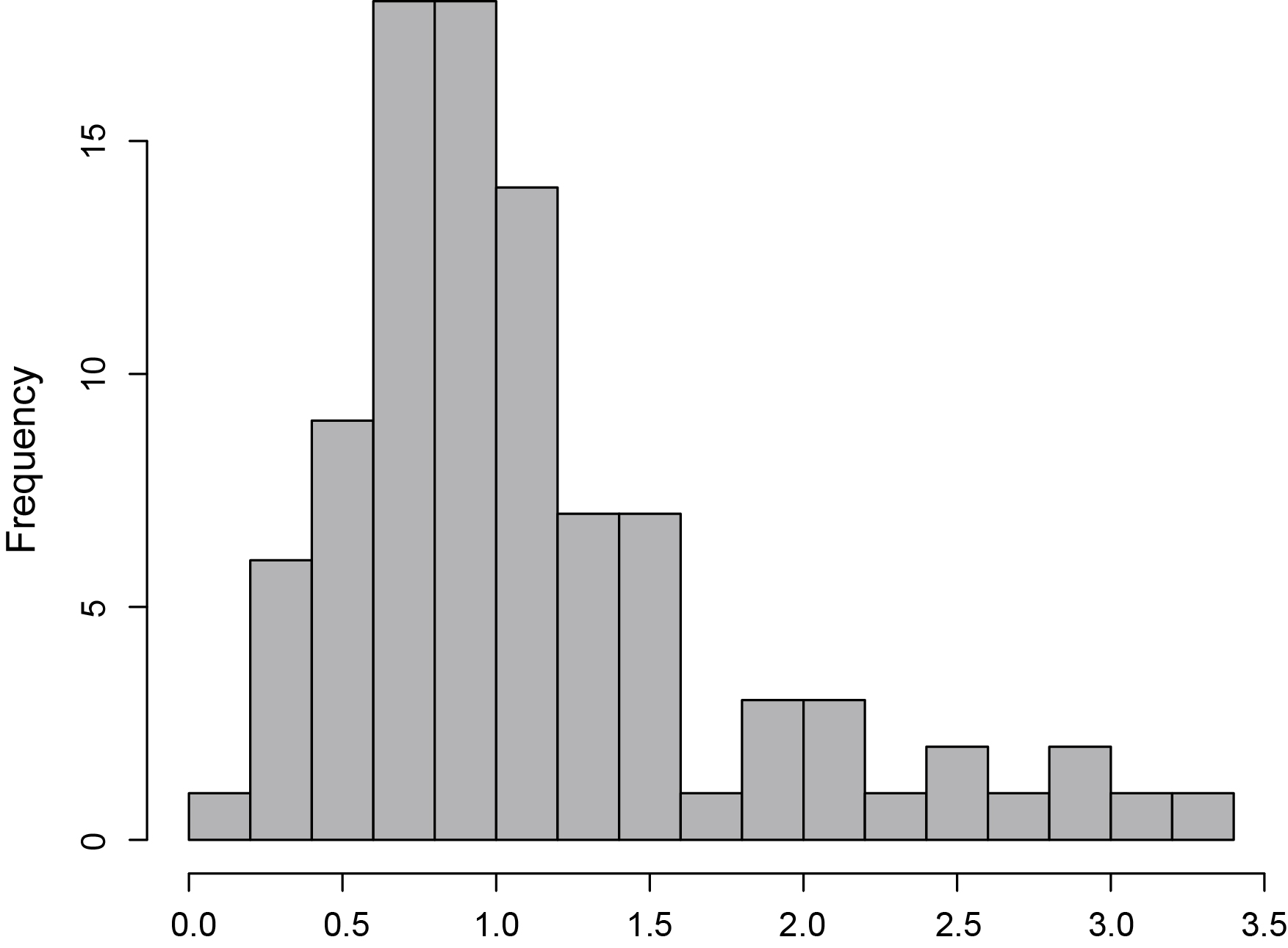

**Supplementary Figure 10.** Determine the threshold of the *Wassertein* distance from housekeeping genes.

The histogram of the largest *Wasserstein* distances between every two clusters for selected housekeeping genes using six mouse datasets (scRNA-Seq: pancreatic islets,

liver, skin; snRNA-Seq: mammary gland(Virgin), skeletal muscle, and epididymal white adipose tissue).
